# A tomato fruit blotch viral replicon defines minimal requirements for cell autonomous replication and identifies functional RNA4-encoded movement and silencing suppression activities

**DOI:** 10.64898/2026.05.21.726790

**Authors:** Niccolò Miotti, Federica Bono, Giulia Golzi, Claudio Ratti, Paola Casati, Massimo Turina, Marina Ciuffo

## Abstract

Tomato fruit blotch virus (ToFBV) is an emerging multipartite positive-sense RNA virus associated with blotchy symptoms on tomato fruits and classified within the genus *Blunervirus* (family *Kitaviridae*). Despite its increasing agricultural relevance, the study of ToFBV has been hindered by the lack of mechanical transmissibility and the difficulty in reproducing infections under controlled conditions. In this work, we report a preliminary step toward the development of the first infectious agroclone system for ToFBV, based on full-length cDNA copies of its four genomic RNAs. We demonstrate that the cloned viral genome is capable of initiating cell autonomous replication in *Nicotiana benthamiana*, as indicated by the accumulation of negative-sense RNA intermediates in infiltrated tissues. To further validate the system, RNA3 was engineered to express GFP, enabling visualization of infection foci and confirming active viral replication in both *N. benthamiana* and tomato. Functional assays of RNA4-encoded proteins demonstrated that it encodes a movement protein capable of complementing movement-deficient viral vectors and a putative suppressor of post-transcriptional gene silencing (PTGS).

Together, these results establish a versatile reverse genetics platform for ToFBV, providing new insights into the replication and functional organization of blunerviruses and enabling future studies on virus–host interactions, pathogenicity, and control strategies.

## INTRODUCTION

Tomato fruit blotch virus (ToFBV; *Blunervirus solani*) is an emerging plant virus associated with characteristic blotchy symptoms on tomato (*Solanum lycopersicum* L.) fruits. It was first identified in Italy and Australia in 2018 (Ciuffo et al., 2020) although retrospective analyses have confirmed its presence in archived symptomatic tomato samples in Italy dating back to 2012, indicating that the virus may have been circulating undetected for several years prior to its formal characterization. Following its initial identification, ToFBV has since been reported across multiple regions in Europe and South America in both field and greenhouse tomato systems, as well as in archived samples (Nakasu et al., 2022; Rivarez et al., 2022; Beris et al., 2023; Bertin et al., 2025). This broad and rapidly emerging geographic distribution raises concerns about the potential economic impact of ToFBV on tomato production. Based on genome organization and phylogenetic placement, ToFBV has been proposed as a member of the genus *Blunervirus* within the family *Kitaviridae*, a relatively recently established group of multipartite positive-sense single-stranded RNA (ssRNA+) viruses (Quito-Avila et al., 2013). The family *Kitaviridae* comprises three genera —*Blunervirus*, *Cilevirus*, and *Higrevirus*— whose members infect a broad range of dicotyledonous plants and are often associated with chlorotic, necrotic, or blotchy symptoms, particularly affecting leaves and fruits. Intriguingly, the RNA-dependent RNA polymerase (RdRP) encoded by *Kitaviridae* members contains domains that are more closely related to those found in arthropod-infecting viruses than in typical plant viruses (Kitajima et al., 2023), suggesting the possibility of an ancient interkingdom horizontal transfer event between arthropodes and plants.

Family members of the genus *Cilevirus* have recently been successfully engineered into infectious agroclone systems for transient expression in model plants. This has provided a valuable tool for studying the biology of Citrus leprosis virus C (CiLV-C) (Leastro et al., 2024) and has also enabled the transient expression of exogenous proteins (Leastro et al., 2025).

The biological cycle of ToFBV, as with other blunerviruses, remains largely unresolved; even with the recent identification of a vector mediating virus spread, the difficulty in achieving reproducible infections limits the dissection of replication mechanisms and genome-encoded functions. Symptomatology in infected tomato plants includes irregular blotches on the fruit surface that become evident during the ripening of the fruits. This element makes managing infections in the field difficult since the symptoms are recognizable mainly during late stages of maturation. Leaves remain symptomless throughout the growing season. A mite vector has been recently demonstrated to be able to transmit the ToFBV (Bertin et al., 2025): *Aculops lycopersici* (Trombidiformes: Eriophyoidea) can act as a vector in tomato and new symptoms, such as leaf chlorosis and reduced growth, have been described in infected tomato plants in controlled conditions that were part of the transmission experiment. The fact that the vector is an eriophyid mite, which is in itself a tomato pest, has a strong impact on the epidemiology of ToFBV, as climatic conditions greatly influence mite population dynamics. In particular, hot and dry summers have been associated with increased eriophyid abundance, which in turn leads to a higher incidence of virus transmission. This helps explain why ToFBV outbreaks fluctuate and have been inconsistent across different years.

Most critically, ToFBV is not mechanically transmissible (Ciuffo et al., 2020), making it impossible to reproduce infections by traditional sap inoculation methods posing a major limitation for the experimental study of the virus, particularly for formally fulfilling Koch’s postulates and conducting controlled pathogenesis or host range studies. Furthermore, members of the family *Kitaviridae* are known to move cell-to-cell in plants inefficiently, significantly reducing systemic infection in plant hosts; this poses another level of complexity for the engineering of these viruses and the replication of natural infections through infectious clones is particularly challenging.

The inability to transmit ToFBV mechanically and the high yield loss often associated with its infections in tomato fields (Kitajima et al., 2023), highlight the urgent need for alternative approaches to investigate its biology. In this context, the development of biotechnological tools, such as infectious clones, can provide a solution. Infectious clones allow researchers to reconstruct viral infections from cloned complementary DNA (cDNA) copies of the viral genome, typically delivered into plants via mechanical inoculation of synthetic transcripts or *Agrobacterium tumefaciens*-mediated transient expression (Ahlquist & Janda, 1984; Wang et al., 2015; Saad et al., 2021). This approach bypasses the need for natural vectors and enables precise control over infection timing and tissue targeting.

In this study, we report the development of agroclones for ToFBV by assembling full-length cDNA copies of its genome segments under the control of plant-active promoters within *Agrobacterium*-compatible binary vectors. These tools allowed the functional characterization of viral open reading frames, including the validation of the cell to cell movement protein activity and the identification of a putative suppressor of post-transcriptional gene silencing (PTGS). From this initial set of clones, some GFP-expressing derivatives were generated, enabling replication-dependent visualization of infection. Together, these tools represent the first significant advancement in the study of blunerviruses, providing a platform for dissecting viral gene function, host-virus interactions, and possibly susceptibility genes to be used to derive resistant lines.

## MATERIAL & METHODS

### Viral source and RNA extraction

Lyophilized tomato leaf samples infected by ToFBV (T1099, PLAVIT collection) kept at -80°C from previous works (Ciuffo et al., 2020) were used. A total RNA extraction from leaves has been carried out using TRIzol™ Reagent (Invitrogen™) following manufacturer’s instructions. Quality and abundance of extracted RNA was checked with Nanodrop One-c (Thermo Fisher Scientific). The same RNA extraction procedure was used to detect the virus in patch areas of agroinfiltrated leaves.

### Infectious clone assembly strategy

Specific primers matching the 5’ and 3’ region of all sequences of the four ToFBV RNAs were designed (Supplementary Table 1). RT has been performed using SuperScript™ IV Reverse Transcriptase following manufacturer’s instruction with random hexamers on 5µl of total RNA extraction. Then, 1µl of diluted cDNA (1:50) has been used as a template for a PCR with Phusion™ High-Fidelity DNA Polymerase following manufacturer’s instruction (annealing 58°C and 2 min extension).

Cloning of RNA 1 and 2 has been done in two steps, dividing both cDNAs in two fragments: unique restriction sites were identified within the ToFBV genomic cDNAs to enable stepwise assembly of full-length cDNA clones in pJL89 (Lindbo, 2007b).

For RNA1, a unique *Kpn*I site at nucleotide position 2205 was selected. Two overlapping fragments were defined: fragment 1A (nt 1-2250) and fragment 1B (nt 2150-5811). For RNA2, a unique *Bam*HI site at nucleotide position 2657 was used to define fragment 2A (nt 1-2700) and fragment 2B (nt 2600-3643).

To prevent unwanted cleavage during cloning, the unique *Kpn*I (RNA1 construct) and *Bam*HI (RNA2 construct) sites present in pJL89 backbone were removed by restriction digestion, end-filling with Klenow fragment (Promega), and blunt-end re-ligation. Loss of the restriction sites was confirmed by diagnostic restriction analysis.

Primers were designed approximately 50 nt upstream and downstream of the selected restriction sites (Supplementary Table 1). Reverse primers included a *Sma*I site present also in pJL89 upstream of the hepatitis delta ribozyme present in the plasmid. Fragment A of each RNA was amplified using a 5′-phosphorylated forward primer (Supplementary Table 1) and cloned into pJL89 digested with *Stu*I and *Sma*I. Plasmid were ligated using T4 DNA ligase (Thermo Fisher Scientific) and then transformed into competent *Escherichia coli* cells. Colony PCR and diagnostic restriction analyses were used to confirm insertion.

Fragment B was amplified and digested with *Kpn*I-*Sma*I (RNA1) or *Bam*HI-*Sma*I (RNA2) and ligated into the corresponding intermediate constructs (pJL89_RNA1A and RNA2A) digested with the same enzymes to reconstitute full-length viral cDNAs.

Cloning of RNA 3 and 4 was done using a pJL89 binary vector linearized with *Stu*I and *Sma*I (Anza Restriction Enzymes, Thermo Fisher Scientific) following manufacturer’s instruction. Dephosphorylation by alkaline phosphatase (Thermo Fisher Scientific) was performed on the pJL89 cut with *Stu*I and *Sma*I, which was then used for cloning the two phosphorylated cDNAs in two distinct reactions with T4 DNA ligase (Thermo Fisher Scientific).

All four plasmid containing ToFBV genome fragments (here after pNM : RNA1/2/3/4) were then used singularly for the transformation of chemically competent DH5α *E. coli* cells. Transformed cells were then plated on selective LB agar added with 50µg/ml of kanamycin and grown at 37°C overnight. Selection of positively transformed colonies has been done using primer pair pJL89_F and a reverse primer specific for RNA 1, 2, 3 and 4 (Supplementary Table 1). Five µl of boiled colonies were used as template for a PCR amplification done with DreamTaq DNA Polymerase (Thermo Fisher Scientific) and presence of expected amplicons were checked on an agarose gel. Overnight growth of screened colonies in LB added with selective antibiotic was performed and plasmid were then isolated with ZymoPURE Plasmid Miniprep Kit following manufacturer’s instructions.

One µg of plasmid was used for the transformation of competent C58C1 *Agrobacterium tumefaciens* strain through heat shock. Transformed cells were then plated on selective LBA petri dishes (kanamycin and tetracycline 50µg/ml) and grown for 2 days.

### ToFBV RNA3 GFP expressing vector

Mutants expressing the green fluorescent protein (GFP) were generated by replacing a portion of ORF2 (nt 756–1532) and ORF5 (nt 2191–2766) of RNA3 with the GFP coding sequence, inserted in-frame with ORF2 or ORF5, respectively, while retaining the first few amino acids of each ORF. Inverted PCR to linearize the pNM : RNA3 plasmid has been performed with Phusion™ High-Fidelity DNA Polymerase following manufacturer’s instruction (55°C annealing and 4:30 min extension). Primer pairs used are reported in Supplementary Table 1; the forward primer has been designed to add *Not*I restriction site. The linearized plasmid was excised from the gel, purified, and subsequently digested with *Not*I, resulting in a linear pNM: RNA3 vector containing a blunt end and a *Not*I site for the in-frame insertion of GFP: previously constructed pJL22 plasmid carrying the GFP sequence was cut with *Stu*I and *Not*I (Anza Restriction Enzymes, Thermo Fisher Scientific) and the product ran on a 1% agarose gel and purified as reported above. Ligation of the linearized plasmid with the GFP insert was performed using T4 DNA ligase (Thermo Fisher Scientific), generating the constructs pNM : RNA3_GFP_ORF2 and pNM : RNA3_GFP_ORF5. Newly generated plasmids were verified by Sanger sequencing to confirm correct cloning. The resulting constructs were then introduced into *A. tumefaciens* C58C1 competent cells as described above.

### ToFBV RNA4 ORF binary expressing vector

Primers for the amplification of RNA4-ORF 1, RNA4-ORF2 and RNA4 1+2 were designed (Supplementary Table 1) adding *Stu*I at 5’ of the forward primers and *Not*I at 5’ of the reverse primers. PCR with Phusion™ High-Fidelity DNA Polymerase was performed using pNM : RNA4 as template, amplicon sizes were checked and then excised from a 1% agarose gel. Purified amplicons were then cloned in Zero Blunt™ PCR (Thermo Fisher Scientific) and used for the transformation of chemically competent DH5α cells. Colonies were screened using M13 and specific reverse primers for ORF1 or 2 (Supplementary Table 1). Plasmids from overnight minipreps of selected colonies were purified and cut with *Stu*I and *NotI*, run on an agarose gel, excised and then ligated into a pJL22 binary vector (Lindbo, 2007a) digested with the same restriction enzymes. Transformation of chemically competent DH5α cells was followed by colony screening using pJL89_F and specific reverse primers for ORF1 or 2 (Supplementary Table 1).

Agroclones developed and used in this study:

- C58C1 pNM :: RNA1
- C58C1 pNM :: RNA2
- C58C1 pNM :: RNA3
- C58C1 pNM :: RNA4
- C58C1 pNM :: RNA3_GFP_ORF2
- C58C1 pNM :: RNA3_GFP_ORF5
- C58C1 pNM :: RNA4_ORF1
- C58C1 pNM :: RNA4_ORF2
- C58C1 pNM :: RNA4_ORF1+2

Detailed structural designs of the C58C1-compatible expression constructs (pNM : RNA3_GFP_ORF2, pNM : RNA3_GFP_ORF5, pNM : RNA4_ORF1, pNM : RNA4_ORF2, and pNM : RNA4_ORF1+2) are illustrated in Figure 1.

**Figure 1:**
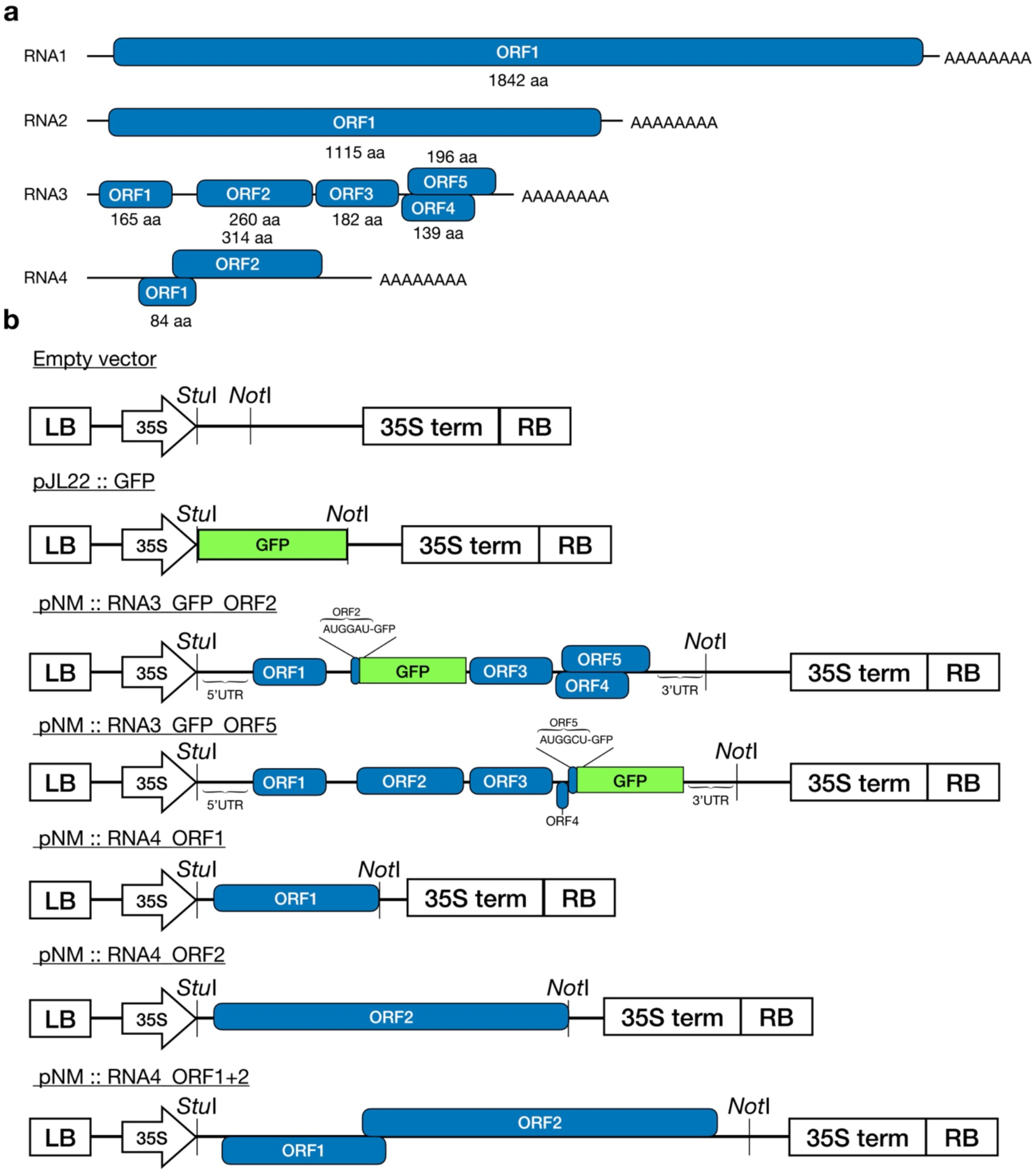
(A) Schematic representation of the four single-stranded RNA segments (RNA1– RNA4) constituting the genome of tomato fruit blotch virus (ToFBV). Blue boxes represent the open reading frames (ORFs) with their corresponding translation products indicated in amino acids (aa). (B) Schematic Representation of the developed recombinant plasmids. Empty vector: The parental binary vector backbone containing the Left Border (LB), the double Cauliflower mosaic virus 35S promoter (35S), the 35S terminator (35S term), and the Right Border (RB). The insertion site features the restriction sites *Stu*I (positioned directly downstream of the 35S promoter) and *Not*I; pJL22 :: GFP construct expressing the green fluorescent protein (GFP) inserted between the *Stu*I and *Not*I restriction sites; pNM :: RNA3_GFP_ORF2 and ORF5 with with modified expression cassettes where ORF2 and ORF5 are substituted by the GFP gene. The first three N-terminal triplets (codons; AUGGAU and AUGGCU) are conserved to keep the GFP gene in the correct reading frame; pNM :: RNA4_ORF1, pNM :: RNA4_ORF2, and pNM :: RNA4_ORF1+2 are the recombinant constructs engineered to express RNA4 components between the *Stu*I and *Not*I restriction sites under the control of the 35S promoter.

### Sequencing of cloned fragments of ToFBV with MinION

Cloned pNM : RNA1,2,3 and 4 of ToFBV were linearized with *Not*I and subjected to full-length sequencing with MinION Mk1B sequencer (Oxford Nanopore Technologies, Oxford, UK) following the manufacturer’s protocol for ligation-based library preparation. Briefly, linearized DNA was end-repaired, dA-tailed, and adapter-ligated prior to loading onto FLO-MIN114 flow cells, according to the instructions provided with the Ligation Sequencing Kit (SQK- NBD114.96). Raw MinION reads were processed using the Galaxy platform. Adapter trimming was performed with Porechop, and reads were subsequently mapped to the reference sequences of ToFBV (MK517477.2, MK517478.2, MK517479.2 and MK517480.2) using Minimap2. Consensus sequences were subsequently assembled from the mapped reads to reconstruct the full-length plasmid inserts and confirm the integrity of the cloned sequences.

### Plant material and agroinfiltration

Model plant *Nicotiana benthamiana* was used for transient expression experiments in this study. Transgenic lines of *N. benthamiana* expressing GFP (16C) have also been used. For tomato transient expression experiments, cv. Marmande has been used. All plants were grown in a growth chamber under controlled conditions, including a 16/8-hour day/night photoperiod, of 25°C and 18°C day/night temperatures, and 70% relative humidity. Three-week-old plants were agroinfiltrated with different combinations of agrobacterium suspensions, including an agrobacterium for the expression of the p19 of tomato bushy stunt virus as a post-transcriptional gene silencing suppressor (Turina et al., 2002). Considered infectious agroclones were grown o.n. in LB added with kanamycin and tetracycline 50µg/ml and MES (2-(N-morpholino) ethanesulfonic acid) 10 mM at 28°C. After pelleting the cultures at 4,500 rpm for 10 min, the cells were resuspended in infiltration buffer (10 mM MES, 10 mM MgCl₂, and 200 µM acetosyringone) to a final OD₆₀₀ of 0.6. Agroinfiltration has been performed using a needless syringe on fully developed leaves of both *N. benthamiana* and tomato cv Marmande. Plants were then maintained in the growth chamber until sampling, which was carried out at different time points ranging from 3–5 days post-inoculation up to 15–30 days post-inoculation, depending on the experimental conditions. Sampled leaves were then conserved at -80°C until further processing. *A. tumefaciens* expressing mCherry under the 35S promoter were a generous gift of Prof. Liying Sun from Northwest A & F University, Yangling, China.

### Detection of ToFBV cDNA and Agroclone-Derived Plasmid DNA

Total nucleic acids were extracted using TRIzol reagent, as previously reported. and used as template for either direct PCR with ToFBV-specific primers (Supplementary Table 1) or for reverse transcription-PCR (RT-PCR) assays. First-strand cDNA synthesis was performed using M-MLV reverse transcriptase (Promega) with random hexamer primers, following the manufacturer’s instructions. Subsequent PCR amplification was carried out using GoTaq Flexi DNA Polymerase (Promega) according to the manufacturer’s protocol. Amplicon presence and size were verified by agarose gel electrophoresis.

For DNA detection, PCR analyses were performed to validate agroinfiltration-based transient transformation and to assess the presence of plasmid DNA within infiltrated patch areas. Prior to RNA-based analyses, DNase treatment was performed as described above to eliminate residual DNA and ensure specific detection of RNA-derived transcripts. Systemic leaves of both *N. benthamiana* and tomato plants were sampled for detection of systemic ToFBV spread, for which only RNA-based detection assays were conducted.

### Validation of active replication via tagged RT-qPCR

A strand-specific RT-tag qPCR was performed to selectively detect the negative-sense ssRNA of ToFBV. Primers were designed to specifically target ToFBV ssRNA1(-) (Supplementary Table 1). A non-viral tag sequence was added to the 5′ end of the forward primer after verifying that it shared no homology with any ToFBV genomic regions, the backbone sequences of the binary vectors used in this study or genomic regions of *N. benthamiana*.

Total RNA was extracted from agroinfiltrated patch areas as described above and 10 µL were treated with DNase I (Invitrogen) to remove residual DNA. Subsequently, 5 µL of DNase-treated RNA were reverse-transcribed using M-MLV Reverse Transcriptase (Thermo Fisher Scientific), according to manufacturer’s instructions, using the forward tagged strand-specific primers (Supplementary Table 1). To eliminate residual tagged primers, cDNA was purified using the DNA Clean & Concentrator kit (Zymo Research).

Quantitative PCR was performed using the tag-specific forward primer (Tag_F), a ToFBV RNA1 specific reverse primer, and a hydrolysis fluorescent probe (Supplementary Table 1) with iTaq Universal Probes Supermix (Bio-Rad), following the manufacturer’s recommendations.

Negative-strand RNA accumulation was normalized to the reference gene *N. benthamiana* F-BOX (Liu et al., 2012., Supplementary Table 1). cDNA was synthesized from DNase-treated RNA using random hexamers and M-MLV Reverse Transcriptase, and qPCR was performed with SYBR™ Green Universal Master Mix (Thermo Fisher Scientific). Relative expression levels were calculated using the ΔΔCt method, and log-transformed ΔΔCt values were plotted.

### Complementation assays of movement-deficient PVX and ToANV GFP-expressing vectors

Previously described movement-deficient mutants of potato virus X (Sun et al., 2013) and tomato apex necrosis virus (ToANV) (Ferriol et al., 2017), lacking the movement protein (MP) open reading frame and expressing GFP, were used in movement complementation assays to evaluate the activity of ToFBV RNA4. The movement protein of tobacco mosaic virus (TMV), previously cloned into a binary expression vector, was used as a positive control for comparison of complementation efficiency. All these clones were described in Bono et al. (2026).

PVX or ToANV agroclones were grown overnight as described above. Agroinfiltration mixtures were prepared by combining the viral constructs with ⅕ serial dilutions of Agrobacterium suspension carrying pNM : RNA4_ORF1 or ORF1+2 ranging from 1:5 to 1:625, in order to obtain predominantly single-cell transient expression events in *N. benthamiana* leaves. Leaf patches were excised at 3 to 5 dpi and GFP fluorescence was evaluated in vivo through confocal microscopy (see details below). Infection foci were identified by GFP fluorescence. GFP-positive infection sites were counted and classified in three classes according to the number of fluorescent cells per focus (single cell, 2–3 cells, and >3 cells), allowing evaluation of virus cell-to-cell movement.

### Confocal microscopy

Leaf samples from agroinfiltrated patch areas were excised using a scalpel, mounted on glass microscope slides, and analyzed by confocal laser scanning microscopy. Samples were analyzed using a Zeiss LSM 880 confocal laser scanning microscope (Carl Zeiss AG, Oberkochen, Germany) equipped with a Plan-Apochromat 20× objective. GFP fluorescence was excited with the 488 nm line of an argon laser and detected within an emission window of 500–525 nm; mCherry was detected using a 600-625 emission window. Chlorophyll autofluorescence was excited using the same 488 nm laser line, with emission collected between 650 and 700 nm. Brightfield images were acquired using the ESID detector. Image acquisition and processing were performed using ZEN 2.3 software (Carl Zeiss AG, Oberkochen, Germany) (Bono et al., 2026).

### In silico structural analysis of ORFs products encoded by the ToFBV genome

For each of the proteins putatively encoded by the ORFs present in RNA 3 and RNA 4 we derived a structural prediction in silico through Alphafold 3 (Abramson et al., 2024). The PDB (Burley et al., 2018), BFVD (Kim et al., 2024) and AlphaFold Protein Structure (Varadi et al., 2023) databases were searched with the most likely model fold as query using FoldSeek (van Kempen et al., 2023). The best matches with some statistical support were displayed with ChimeraX (Meng et al., 2023).

## RESULTS AND DISCUSSION

### Transiently expressed ToFBV fragments accumulates ssRNA1(-) in patch areas of *N. benthamiana*

The ToFBV genome consists of four RNA segments (Figure 1). The first two RNAs are monocistronic and are putatively associated with viral replication (MK517477.2 and MK517478.2). RNA1 encodes a large multifunctional replication-associated protein containing conserved domains typical of positive-sense RNA viruses, including an alphavirus-like methyltransferase domain, an OTU-like cysteine protease domain, a P-loop NTPase/helicase domain, and a (+) RNA virus helicase core domain. Notably, OTU-like proteases identified in several RNA viruses have also been associated with deubiquitinating activities involved in suppression or modulation of host antiviral immune responses (James et al., 2010; van Bailey-Elkin et al., 2014), suggesting a potential additional role of the ToFBV RNA1-encoded protein in host–virus interactions. In contrast, RNA2 encodes the RNA-dependent RNA polymerase (RdRP), which provides the catalytic activity required for viral RNA synthesis (Ramos-González et al., 2023). This genomic organization is consistent with members of the genus *Blunervirus*, in which the replication machinery is functionally divided between RNA1, encoding methyltransferase-, protease-, and helicase-related activities, and RNA2, encoding the RdRP.

MINION sequencing confirmed that the cloned pNM::RNA1, RNA2, RNA3, and RNA4 constructs were highly consistent with the reference sequences previously isolated and characterized from the T1099-infected plant. Pairwise sequence alignments showed approximately 99% nucleotide identity compared with the reference sequences. All identified mutations were further analyzed and classified as synonymous or nonsynonymous (Supplementary Table 2).

To assess the ability of the developed agroclones to initiate viral replication in agroinfiltrated leaves, different mixtures of agrobacterial suspensions were tested. A complete infection containing all four ToFBV genomic fragments (hereafter “ToFBV complete”) was compared with an agroinfiltration lacking the RNA2 construct (ΔRNA2), that codes for the viral RdRP. An additional mix of agroclones consisted of the agroinfiltration of RNA1 and RNA2 expressing clones only, to evaluate whether replication could occur in the absence of RNA3 and RNA4 (hereafter “RNA1+2”).

Because negative-sense RNA molecules act as replication intermediates during the life cycle of positive-sense RNA viruses, their detection is considered evidence of active viral replication (Weber et al., 2006). In this study, the quantification of ssRNA1(-) was used as a proxy to compare the accumulation of replication intermediates generated by the viral RdRP in ToFBV complete infections.

At 5 dpi, *N. benthamiana* leaves showed localized discoloration in patch areas and a darker green rim in the ToFBV complete treatment, whereas no visible alterations in pigmentation or morphology were observed in the ΔRNA2 treatment (Supplementary Figure 1). A similar discoloration was also observed in patch areas agroinfiltrated with RNA1+2 (data not shown). In contrast, tomato plants also developed symptoms, including chlorosis and alterations in leaf morphology; however, these symptoms were observed in both the complete ToFBV and ΔRNA2 experiments.

Quantification of ssRNA1(-) was performed in *N. benthamiana* samples at 5 dpi using an RT-tag qPCR protocol. Accumulation of ssRNA1(−) was up to eight-fold lower in ΔRNA2 samples compared to the complete mix of ToFBV clones (Supplementary Figure 2), supporting the role of the viral RdRP (RNA2) in local replication within infiltrated patches. Interestingly, ssRNA1(−) accumulation in the RNA1+2 treatment was comparable to, or even higher than, that observed following complete ToFBV agroinfiltration, with a mean relative expression (log fold) value of 1 (Supplementary Figure 2). These results suggest that the RNA1 and RNA2 viral fragments alone are sufficient to support ssRNA1(−) accumulation in the absence of the ORFs encoded by RNA3 and RNA4. The residual accumulation of ssRNA1(−) observed in the ΔRNA2 samples may instead be attributed to unspecific transient expression of the RNA1 transcript delivered by agroinfiltration, rather than to bona fide RNA-dependent replication.

The same experimental approach was applied to tomato plants, and agroinfiltrated leaf patches were collected at 5 dpi. We compared ToFBV complete mix, to the ΔRNA2. Also in this case, evidence of RNA2-dependent accumulation was gathered, since the complete mix reached the exponential curve circa 3 Ct earlier than the Δ2RNA mix (Supplementary Table 3). This was confirmed through both RNA1 and RNA4 detection. Regarding detection of the ssRNA(-) of RNA 1, again we showed a difference between the ToFBV complete agroclone set and the ΔRNA2 combination. These results are consistent with those obtained in *N. benthamiana* and confirm that RNA2 is essential for the accumulation of ssRNA1(-) during local infection also in tomato (Supplementary Table 3). Molecular detection of DNA in total nucleic acid extracts from patch areas confirmed the presence of the cloned ToFBV DNA fragments in both hosts, indicating successful delivery of all the constructs as designed.

At 10 dpi, RT-qPCR analysis of systemic leaves of *N. benthamiana* revealed substantial viral RNA accumulation, with Ct values of 19 for RNA4, detected by SYBR Green, and 27 for RNA1, detected by TaqMan (Supplementary Table 3). Systemic leaves from 14 plants of *N. benthamiana* were collected and analyzed at 15 dpi. All ToFBV genomic RNAs (RNAs 1, 2, 3, and 4) were detected in 4 of the 14 samples, while RNA1 was still detectable in 11 out of 14 samples (Supplementary Table 3). In contrast, all systemic samples tested negative for ToFBV DNA fragments, demonstrating the absence of Agrobacterium-derived plasmid DNA. As expected, the local agroinfiltrated leaf areas remained positive for ToFBV DNA, confirming the persistence of the delivered T-DNA at the infiltration sites; we also tested four plants that were inoculated with a ΔRNA2 combination, and they were all negative.

Overall, these results indicate that the developed agroclones efficiently initiate local infection and viral replication in both *N. benthamiana* and tomato, as demonstrated by the accumulation of viral ssRNA(-) and by the RNA2-dependent higher accumulation of viral RNAs. Notably, the constructs also supported systemic infection in *N. benthamiana*, indicating that transient expression of the viral cDNAs was sufficient to generate infectious viral RNAs capable of long-distance movement. The temporal reduction in viral RNA accumulation may reflect the non-host status of *N. benthamiana* for ToFBV, which could limit sustained viral replication and systemic infection. Although members of the *Kitaviridae* family are generally characterized by relatively slow systemic movement, the observed decrease in viral RNA accumulation at later time points suggests that host compatibility may also play an important role in determining viral accumulation in systemic leaves of *N. benthamiana*.

### Engineering of a ToFBV-based GFP-expressing viral vector

To further validate local replication of the cloned ToFBV genome and assess its ability to express a heterologous protein, RNA3 was engineered to express GFP. ORF2 or ORF5 were individually disrupted by partial deletion using inverse PCR, leaving only short N-terminal protein residues, and replaced with the GFP coding sequence fused in frame, generating two constructs (hereafter GFP2 and GFP5). Based on previous results indicating that RNA3 and RNA4 are not essential for RNA1- and RNA2-mediated replication, these constructs were also used to evaluate which genomic context is more suitable for heterologous protein expression of GFP.

Agroinfiltration of *N. benthamiana* leaves was performed with either pNM :: RNA3_GFP_ORF2 (GFP2) or pNM :: RNA3_GFP_ORF5 (GFP5). ToFBV complete and ΔRNA2 were included as controls. All agroinfiltrated plants displayed localized discoloration in infiltrated areas, with the exception of the ΔRNA2 treatment, which showed no visible alterations. Leaf patch areas were collected at 5 days post-inoculation (dpi) and analyzed by confocal microscopy. No fluorescence signal was detected in the ΔRNA2 control under GFP-specific settings demonstrating that no GFP-specific fluorescence was derived from internal initiation of translation of the 35S derived RNA3 expression. In contrast, discrete foci of fluorescent cells were clearly detected in both GFP2 and GFP5 treatments that included both RNA1 and RNA2 expressing agrobacterium suspensions when imaged under GFP-specific conditions (Figure 2), confirming expression of the heterologous protein and, indirectly, active viral replication (because it is RNA2-dependent). The GFP signal was predominantly localized at the cell membrane and within the nucleus, consistent with accumulation of GFP in transiently expressing cells. In the corresponding ΔRNA2 controls, weak fluorescent signal was observed, indicating non-specific fluorescence. This signal was not consistent with GFP expression derived from viral replication, as no defined subcellular localization (e.g., nuclear or plasma-membrane-associated pattern) was detectable (Figure 2). Qualitative differences were observed between the two constructs: GFP5 consistently showed a higher fluorescence intensity compared to GFP2 (Figure 2, Mann–Whitney U test, p < 0.001, with a large effect size r = 1). These results indicate that disruption of ORF5 provides a more favourable genomic context for heterologous protein expression than disruption of ORF2, possibly due to reduced interference with essential or structurally constrained viral functions, or to increased accumulation of subgenomic RNAs in the ORF5 context enhancing GFP expression.

**Figure 2:**
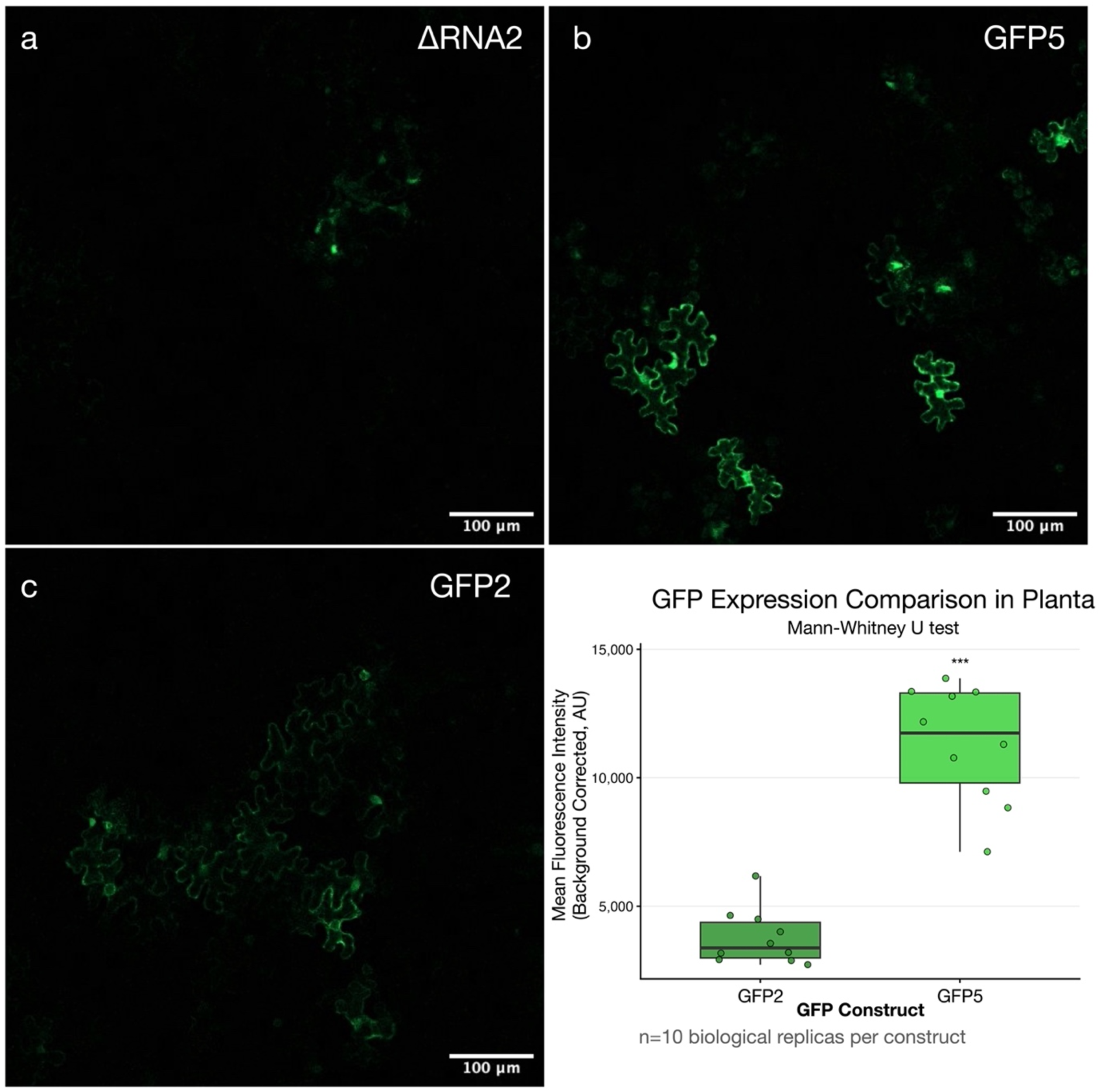
Evaluation of GFP intensity with different ToFBV viral vector constructs in model plants. (A–C) Representative confocal microscopy images of *N. benthamiana* epidermal cells: (A) Negative control (ΔRNA2) showing no detectable specific fluorescence in the absence of the viral replicase, (B) GFP5 construct exhibiting high-intensity specific fluorescence outlining the cell periphery and cytoplasm, (C) GFP2 construct showing successful but visibly lower fluorescence intensity compared to GFP5. Scale bars = 100 µm. (Bottom Right) Quantitative analysis of Mean Fluorescence Intensity (Background Corrected, AU). The box plot displays the distribution of 10 biological replicates per construct, with individual data points overlaid. The median intensity of GFP5 is significantly higher than that of GFP2 (Mann-Whitney U test, p < 0.001).

In parallel, agroinfiltration experiments with GFP5 were conducted in tomato plants, the natural host of ToFBV. Confocal microscopy analysis revealed discrete foci of GFP-positive cells in infiltrated areas after 5 dpi (Figure 3), demonstrating that local replication of the engineered virus is achievable in tomato. In ΔRNA2 controls, lacking the expression of RNA2, it was not possible to identify any fluorescence determined by accumulation of expressed GFP (Figure 3). These results provide, to our knowledge, the first direct evidence of replication of an engineered *Blunervirus* in its natural host using an agroclone-based system.

**Figure 3:**
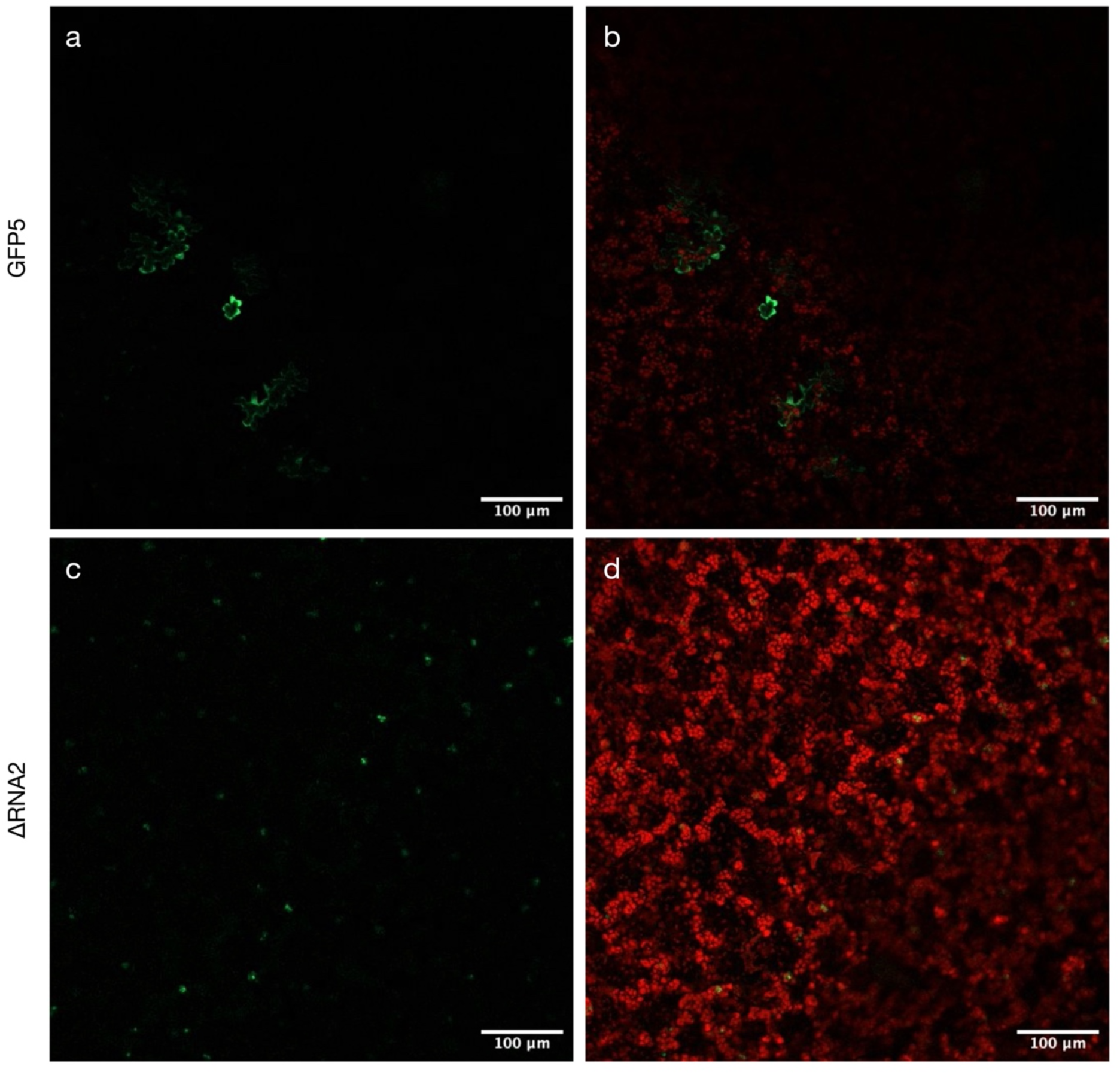
Validation of viral vector-mediated GFP expression in tomato (S. lycopersicum cv. Marmande) leaf tissue. (A, B) Representative confocal microscopy images of leaf tissue infected with the GFP5 construct. (A) Green channel showing robust GFP expression in epidermal cells. (B) Merged image of GFP (green) and plant chlorophyll autofluorescence (red), illustrating the distribution of the viral vector relative to the underlying mesophyll structure. (C, D) Negative control images of leaf tissue treated with the ΔRNA2 mix constructs. (C) Green channel demonstrating the absence of detectable specific GFP fluorescence when the viral replicase is missing (some auto fluorescent spots are present but do not match cellular shape. (D) Merged image showing the red chlorophyll auto fluorescence, confirming that the faint signals in (C) represent unspecific fluorescence rather than GFP-specific protein expression. Scale bars = 100 µm.

Transient expression assays performed with the ToFBV GFP-expressing viral vector revealed clear differences in GFP accumulation and cell-to-cell spread between *N. benthamiana* and tomato plants (Figure 2 and 3). In both hosts, GFP fluorescence was successfully detected, confirming that the viral constructs were functional and capable of initiating infection. However, fluorescent foci observed in *N. benthamiana* were consistently larger and involved a greater number of adjacent cells compared to those detected in tomato, where fluorescence was generally restricted to smaller and more isolated cell clusters. This difference is likely associated with the lower efficiency of Agrobacterium-mediated co-delivery in tomato tissues. Since the establishment of a productive ToFBV infection requires the simultaneous delivery of all four viral RNA segments into the same cell, reduced transient co-expression efficiency may represent a major limiting factor in tomato. To investigate this hypothesis, co-expression experiments using GFP- and mCherry-expressing constructs were performed in both hosts. In *N. benthamiana*, a high proportion of cells simultaneously expressing both fluorescent proteins were observed, indicating efficient co-expression. In contrast, tomato tissues predominantly displayed cells expressing either GFP or mCherry alone, with rare double-fluorescent cells (Supplementary Figure 3). These observations support the hypothesis that the reduced efficiency of four simultaneous agroclone delivery in tomato substantially limits the establishment and spread of complete ToFBV infections in its natural host.

### AlphaFold-based structural analysis of RNA3 and RNA4 ORFan proteins

RNA3 of ToFBV (MK517479.2) contains five predicted ORFs, all encoding proteins lacking significant sequence similarity to entries in publicly available databases, and thus classified as ORFans (Ramos-González et al., 2023). Among these, ORF2 and ORF3 exhibit conserved features shared with other members of the genus *Blunervirus*. In particular, ORF2 contains predicted transmembrane helices (TMHs) near its C-terminus, suggesting a possible association with cellular membranes. ORF3 encodes a homolog of the SP24 protein, which is conserved across members of the family *Kitaviridae* (Ramos-González et al., 2023).

To gain functional insights into these ORFan proteins, structure predictions were generated using AlphaFold and compared against structural databases using Foldseek (see Materials and Methods). Among the five RNA3-encoded proteins, only ORF1 and ORF3 yielded some regions with confident structural predictions (based on pLDDT scores), predominantly consisting of α-helical elements (Figure 4). The remaining ORFs (ORF2, ORF4, and ORF5) showed overall low-confidence predictions, preventing reliable structural interpretation. Foldseek-based structural similarity searches revealed that ORF1 shares structural features with the p14 protein (A0A386QVI9) of tea plant necrotic ring blotch virus, another member of the genus *Blunervirus* (Figure 4). This possible structural conservation suggests a possible conserved functional role, however, as p14 itself is an ORFan protein with poorly characterized function, the biological significance of this similarity remains unclear. ORF3 displayed broader structural similarities, not only to homologs from blueberry necrotic ring blotch virus (M4YSN2; *Blunervirus*), but also to proteins from more distantly related members of the *Kitaviridae* family, including pistachio virus X (A0A7T0M858; *Higrevirus*) and hibiscus yellow blotch virus (A0A890CSA5; *Cilevirus*). This observation is consistent with the conserved nature of SP24-like proteins and supports the hypothesis that ORF3 may play a conserved role within the *Kitaviridae* infection cycle in plants or in the possible insect vectors. Overall, the lack of confident predictions and database matches for ORF1, ORF4, and ORF5 highlights the high level of divergence of these proteins and limits functional inference based solely on in silico approaches.

**Figure 4:**
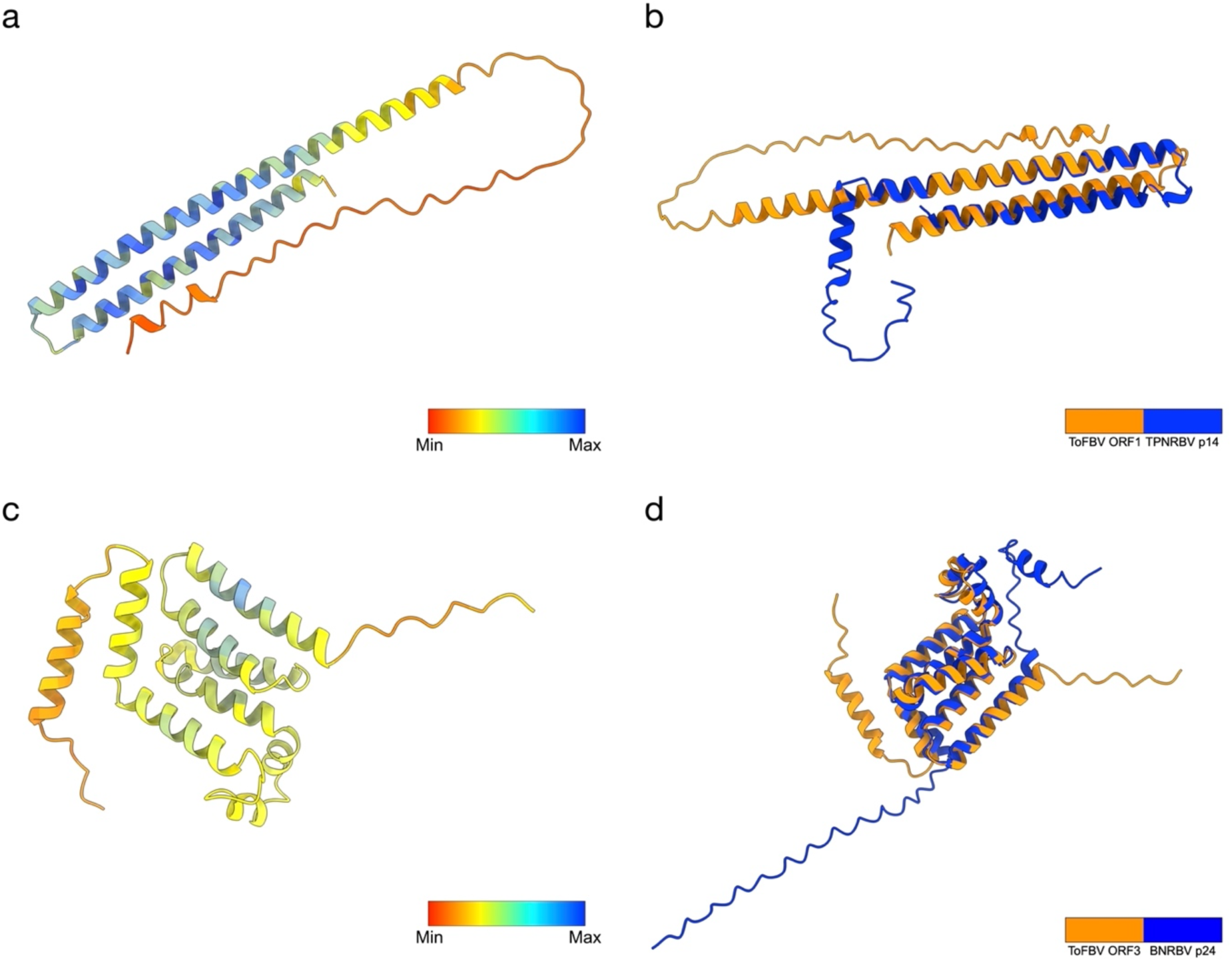
Structural modeling and comparative analysis of ToFBV ORF1 and ORF3 proteins. (A, C) In silico protein structure predictions for ToFBV (A) ORF1 and (C) ORF3 generated using AlphaFold2. The color gradient represents the predicted Local Distance Difference Test (pLDDT) scores. (B) Structural superposition (matchmaking) between the predicted ToFBV ORF1 (orange) and the p14 protein of TPNRBV (blue), showing partial structural conservation. (D) Structural superposition between the predicted ToFBV ORF3 (orange) and the p24 protein of BNRBV (blue), highlighting the folded domains shared between these viral proteins.

RNA4 of ToFBV (MK517480.2) contains two partially overlapping ORFs after a long 5’ UTR. ORF1 begins at nucleotide position 427 and encodes a small protein of 84 amino acids, whereas ORF2 starts at nucleotide position 647 and encodes a 314 amino acid protein. Previous analyses identified a conserved 3A-like motif within ORF2 (Ramos-González et al., 2023), suggesting a potential role in viral movement.

Consistent with this hypothesis, AlphaFold prediction of ORF2 resulted in a well-defined structure with high-confidence regions (Figure 5). Foldseek analysis revealed significant structural similarity to the movement protein of blueberry necrotic ring blotch virus (M4YFU4), as well as to movement proteins from other plant viruses, including tobacco mottle virus (Q9DX75; *Tombusviridae*) and cucumber mosaic virus (O40979; *Bromoviridae*) (Figure 5).

**Figure 5:**
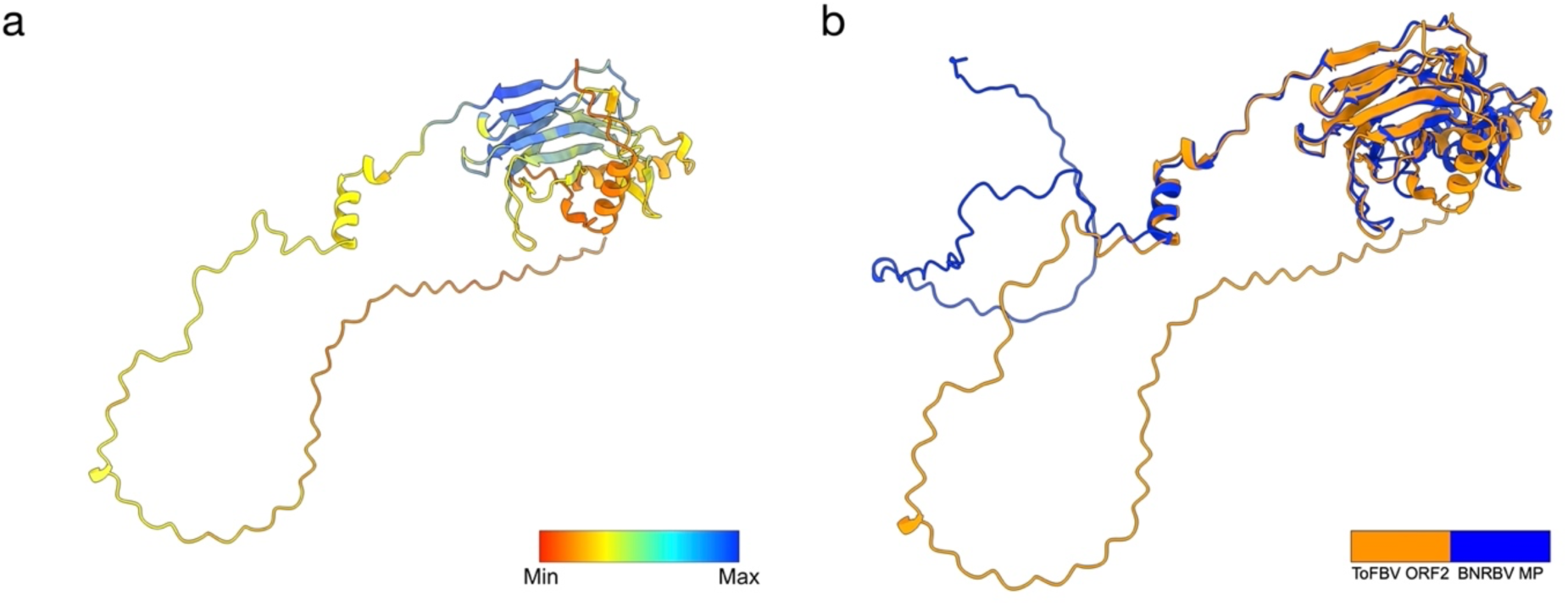
Structural modeling and comparative analysis of the ToFBV ORF2 protein. (A) Three-dimensional structural prediction of the ToFBV ORF2 protein generated via AlphaFold2. Local Distance Difference Test (pLDDT) scores are presented. (B) Structural superposition (matchmaking) between the predicted ToFBV ORF2 protein (orange) and the movement protein (MP) of BNRBV (blue). The alignment reveals significant structural homology, particularly within the globular domain, suggesting a conserved functional role in viral movement.

These findings strongly support the annotation of RNA4 ORF2 as a movement protein and suggest functional conservation despite limited sequence similarity. In contrast, RNA4 ORF1 yielded low-confidence structural predictions, and no significant similarities were identified through Foldseek searches, leaving its potential role unresolved.

A summary of the main Foldseek results in the BFDB database is reported in Table 1:

**Table 1:** Top structural alignment hits for predicted ToFBV proteins within the Big Fantastic Virus Database (BFVD) using Foldseek. Structural similarity metrics (probability, sequence identity, and E-value) are shown for queries from RNA3 and RNA4 against related viral target proteins

| ToFBV | Target | Scientific Name | Probability | Sequence Identity | E-Value |
| --- | --- | --- | --- | --- | --- |
| RNA3 ORF1 | A0A386QVI9 | Tea plant necrotic ring blotch virus | 0.44 | 11.7 | 3.43e-1 |
| RNA3 ORF3 | M4YSN2 | Blueberry necrotic ring blotch virus | 1.00 | 21.4 | 1.83e-8 |
|  | A0A7T0M858 | Pistachio virus X | 1.00 | 20 | 8.59e-7 |
|  | A0A890CSA5 | Hibiscus yellow blotch virus | 1.00 | 18.2 | 7.08e-5 |
| RNA4 ORF2 | M4YFU4 | Blueberry necrotic ring blotch virus | 1.00 | 35.5 | 8.20e-26 |
|  | Q9DX75 | Tobacco mottle virus | 1.00 | 18.7 | 1.74e-13 |
|  | O40979 | Cucumber mosaic virus | 1.00 | 17.9 | 1.84e-13 |

### Functional characterization of ToFBV RNA4: movement protein complementation and PTGS suppressor activity

#### Movement protein complementation assay

To determine whether RNA4 encodes a functional movement protein (MP), complementation assays were performed using movement-deficient GFP-expressing viral vectors derived from PVX and ToANV. Agrobacterium suspensions mixtures were serially diluted to obtain predominantly single-cell infection events; a dilution of 1:625 was found to be optimal for achieving isolated transient expression events, thereby enabling accurate assessment of cell-to-cell movement. Fluorescent infection foci were analyzed at 3 and 5 days post-inoculation (dpi) by confocal microscopy.

Because RNA4 contains two partially overlapping ORFs, complementation assays were initially conducted using a construct expressing the entire RNA4 coding region (ORF1+ORF2). This approach preserves the native genomic architecture and minimizes potential artefacts arising from disruption of overlapping coding sequences, which in RNA viruses are known to impose strong evolutionary constraints due to genome compactness and can influence translational regulation, protein expression, and multifunctional RNA element activity (Ho et al., 2021). The movement protein of tobacco mosaic virus (TMV) was used as a positive control.

At 3 dpi, single-cell infection events were consistently observed in areas infiltrated with the empty vector, confirming the absence of viral movement. In contrast, multicellular fluorescent foci were detected in both the TMV control and the ORF1+ORF2 treatments in the PVX- and ToANV-based systems, indicating successful complementation of cell-to-cell movement. Infection foci frequently comprised more than five cells, demonstrating efficient spread from the initially infected cell. The extent of movement mediated by the ORF1+ORF2 construct appeared comparable to that observed with the TMV MP (Figure 6). Differences in the distribution of the size of infection foci were assessed using Fisher’s exact test. A significant overall effect was observed (p < 0.001) in both PVX (Figure 6) and ToANV systems Supplementary Figure 4). Pairwise comparisons with Benjamini–Hochberg correction showed that the empty vector differed significantly from both TMV and ToFBV (adjusted p < 0.01), whereas no significant difference was observed between TMV and ToFBV (adjusted p > 0.05) (Figure 6). Notably, in the PVX system, patch areas at 10 dpi exhibited fluorescence levels detectable using a portable UV lamp (Supplementary Figure 5). The PVX mutant complemented with ToFBV ORF1+2 showed a significantly larger fluorescent area compared to the TMV MP control.

**Figure 6:**
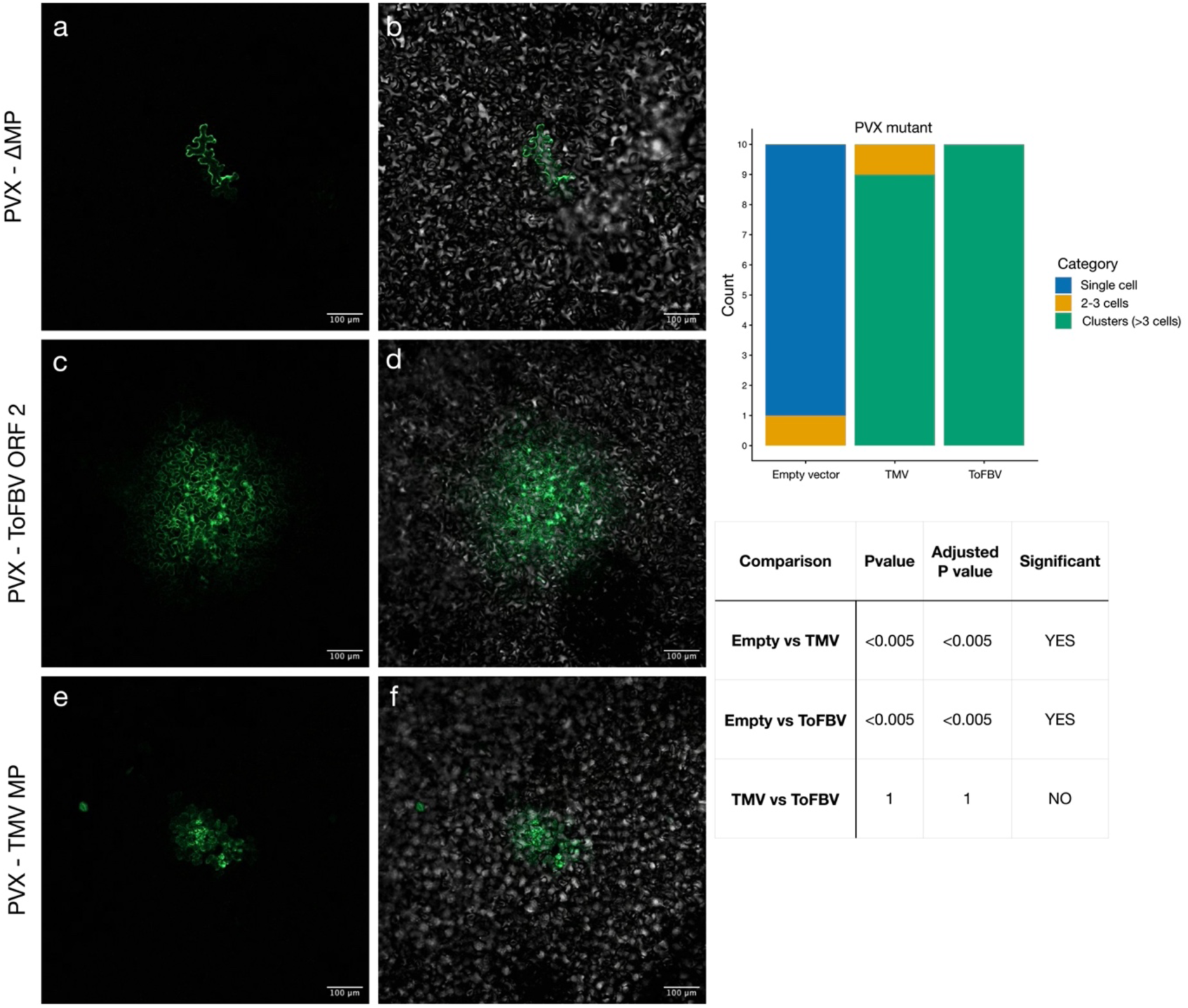
Functional complementation of movement-deficient viral mutants by ToFBV RNA4 ORF2. (A–F) Representative confocal microscopy images of leaf tissue at 3 dpi following a 1:625 dilution infiltration. On the left (A, C, E) show the GFP channel; on the right (B, D, F) show the GFP signal merged with the brightfield channel. (A, B) Negative control: PVX mutant lacking its native movement protein (MP), showing restricted infection limited to single cell. (C, D) Complementation of the PVX mutant by ToFBV RNA4 ORF2, demonstrating restored cell-to-cell movement and the formation of large infection clusters. (E, F) Positive control: complementation by the TMV movement protein, also resulting in multi-cellular cluster formation. Scale bars = 100 µm. (Right) Quantitative analysis of viral spread in both PVX mutant systems. Stacked bar graphs (top and middle) categorize infection foci as single cells, 2–3 cell groups, or clusters (>3 cells) for the empty vector, TMV, and ToFBV constructs. The summary table (bottom) provides statistical comparisons (Adjusted P-values), indicating that ToFBV ORF2 significantly promotes viral spread compared to the negative control (p < 0.005) and performs at a level statistically comparable to the TMV MP.

These results indicate that RNA4 of ToFBV encodes a factor capable of mediating cell-to-cell movement in *N. benthamiana*. In the presence of the ORF1+ORF2 construct, movement-deficient PVX and ToANV mutants regained the ability to spread to adjacent cells, forming multicellular infection foci. Notably, the efficiency of complementation was comparable to, and in some instances appeared to exceed, that observed with the TMV MP under the same experimental conditions.

To further dissect the contribution of individual ORFs, constructs expressing ORF1 or ORF2 alone were tested in the PVX-based system. At 3 dpi, expression of ORF1 alone failed to restore movement, as only single-cell infection events were observed, similar to the negative control. In contrast, ORF2 alone was sufficient to restore cell-to-cell movement, producing multicellular foci at frequencies comparable to those observed with the ORF1+ORF2 construct and the TMV MP control (Supplementary Figure 6). These results strongly suggest that ORF2 encodes the functional movement protein, as anticipated from the results of protein structure prediction presented above, whereas ORF1 does not contribute directly to this activity under the conditions tested.

To investigate the ability of ORF2 to support local spread for ToFBV infection, auto-complementation assays were performed using pNM : RNA3_GFP5 construct. In these experiments, RNA4 was omitted and the movement protein ORF2 was supplied in trans either via pNM :: RNA4_ORF2 or, as a control, an empty vector (pJL22). Serial dilution agroinfiltration was used to favour single-cell infection events and allow the assessment of early local spread. At 6 days post-infiltration, fluorescence microscopy revealed clear cell-to-cell spread of GFP signal in samples where MP was provided in trans (Figure 7). In contrast, in the absence of MP expression, only isolated fluorescent cells were observed, with no formation of multicellular fluorescent clusters (Figure 7), indicating a failure in local viral movement.

**Figure 7:**
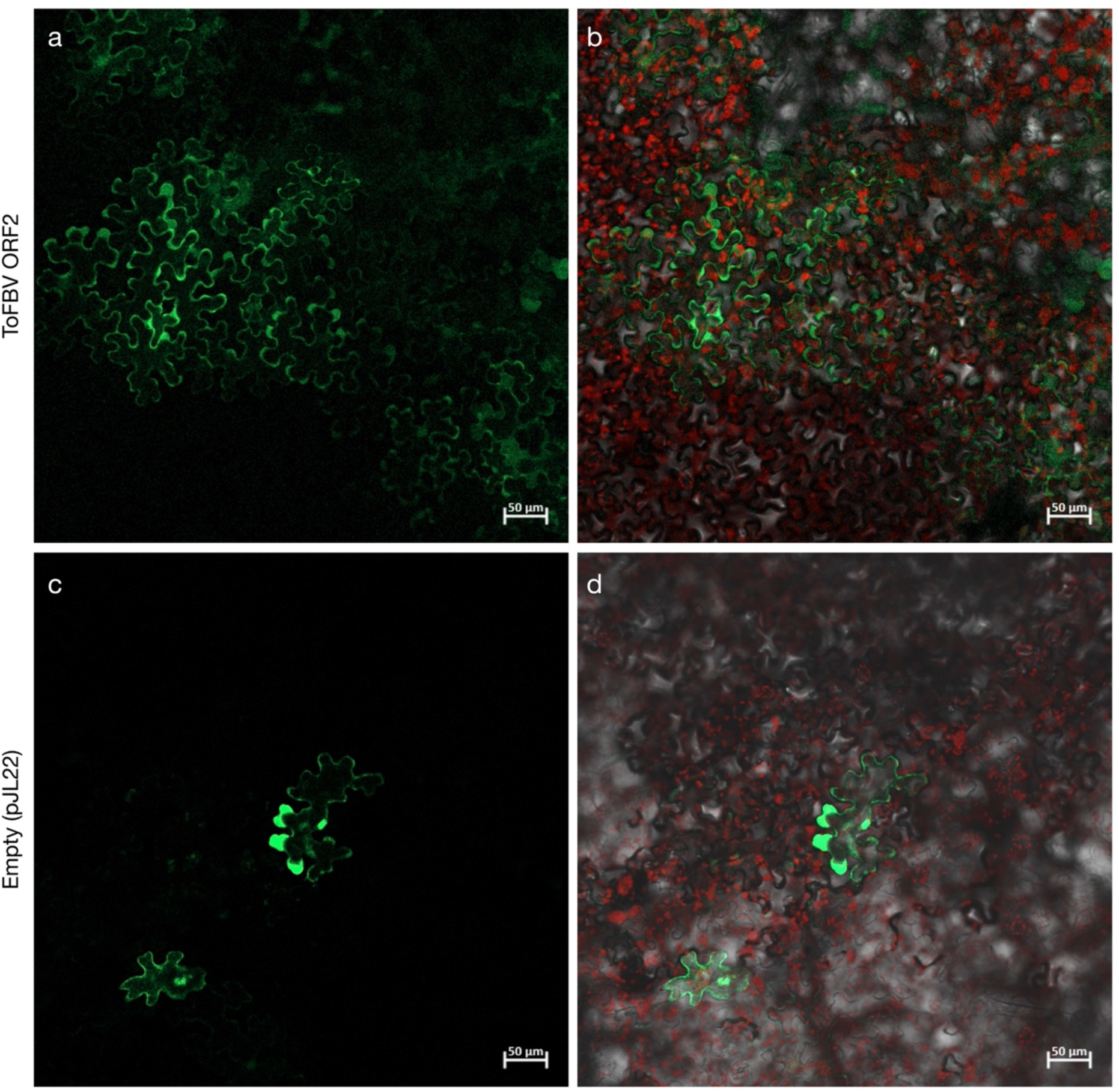
Functional auto-complementation assay of the ToFBV MP. Confocal laser scanning microscopy images of epidermal cells 6 days post-inoculation showing the cell-to-cell movement dynamics of a ToFBV GFP-expressing vector (ToFBV-GFP5). (A, B) Complementation with ToFBV MP (ORF2): (A) GFP fluorescence channel revealing a widespread, multicellular cluster of fluorescent cells, indicating active cell-to-cell viral spread facilitated by the co-expressed MP. (B) Overlay image combining GFP fluorescence, chlorophyll auto fluorescence (red), and brightfield. (C, D) Negative control omitting the MP (pJL22 empty vector): (C) GFP fluorescence channel showing that infection foci remain strictly constrained to single, isolated cells, demonstrating that the virus is unable to spread without the functional MP. (D) Overlay image of the empty vector control confirming the lack of intercellular movement into neighboring tissue. Both experimental conditions were performed using a 1:25 serial dilution to favor single-cell transient expression events. Scale bars = 50 µm.

These results provide the first evidence that transiently expressed ToFBV-derived clones are capable of mediating cell-to-cell movement in a model plant system when the movement protein is supplied.

#### PTGS suppressor assay

Because no canonical RNA silencing suppressor has been identified in the ToFBV genome, we investigated whether RNA4 possesses PTGS suppressor activity. To this end, we employed the well-established GFP silencing assay in transgenic *Nicotiana benthamiana* line 16C, which constitutively expresses GFP.

Leaves of 16C plants were agroinfiltrated with *A. tumefaciens* C58C1 carrying constructs expressing either ORF1 alone (pNM :: RNA4_ORF1), ORF2 alone (pNM :: RNA4_ORF2), or the combined ORF1+ORF2 region (pNM :: RNA4_ORF1+2). In this assay, PTGS suppressor activity is indicated by the retention of GFP fluorescence in infiltrated tissues, whereas loss of fluorescence reflects the onset of GFP silencing. Leaves infiltrated with the well-characterized PTGS suppressor p19 served as a positive control.

Plants were monitored at 4 and 6 days dpi and examined under UV illumination (Figure 8). Co-expression of ORF1 with GFP resulted in detectable fluorescence as early as 4 dpi, suggesting that ORF1 functions as a PTGS suppressor (Figure 8). Interestingly, GFP fluorescence observed in ORF1-infiltrated tissues at 4 dpi appeared stronger than that detected in p19-infiltrated tissues at the same time point. In contrast, the suppressor activity of p19 became clear at 6 dpi (Figure 8). These observations suggest that ORF1 may act as a highly efficient PTGS suppressor, although quantitative analyses will be required to confirm this hypothesis.

**Figure 8.**
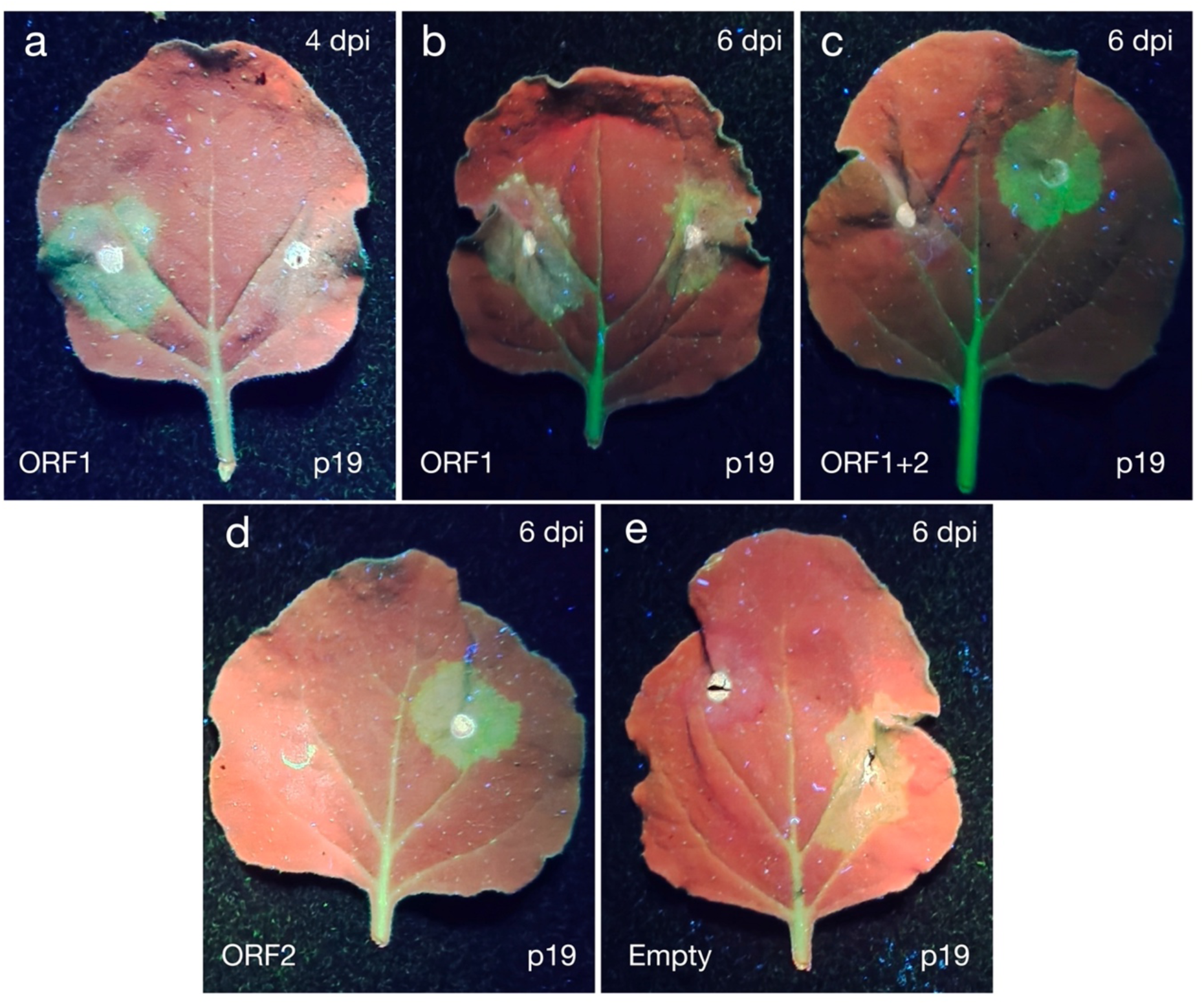
Assessment of PTGS suppressor activity of ToFBV RNA4 and its encoded proteins in N. benthamiana 16C plants. Transgenic *N*. *benthamiana* line 16C, constitutively expressing GFP, was agroinfiltrated with constructs expressing ToFBV individual ORFs of RNA4 to evaluate their ability to suppress PTGS. Leaves were photographed under UV illumination. Green fluorescence indicates suppression of GFP silencing, whereas the absence of fluorescence (dark/red appearance) indicates active silencing of the GFP transgene. In all panels, the right side of the leaf was infiltrated with the positive control p19, while the left side was infiltrated with the indicated test construct. (A) ORF1 retained GFP fluorescence at 4 dpi and 6 dpi (B), suggesting that this protein functions as a PTGS suppressor. Notably, p19-mediated suppression was more evident at 6 dpi than at 4 dpi. In contrast, ORF2 (C) and the combined ORF1+ORF2 construct (D) failed to maintain GFP fluorescence at 6 dpi, indicating a lack of detectable suppressor activity. An empty vector negative control is shown in (E).

In contrast, expression of ORF2 alone or the combined ORF1+ORF2 construct failed to maintain GFP fluorescence, indicating a loss of suppressor activity in these contexts (Figure 8). Since ORF1 partially overlaps with ORF2, the lack of activity observed for the ORF1+ORF2 construct may result from altered expression caused by the absence of the native untranslated regions (UTRs). Notably, the full-length RNA4 construct, which retains both ORFs together with their native UTRs, exhibited suppressor activity. These findings suggest that the proper expression and/or functionality of ORF1 may depend on the genomic context of RNA4, including the presence of its native UTR sequences.

Additional experiments were performed using different combinations of ToFBV genomic components, including the complete viral genome and genome variants lacking either RNA2 (ΔRNA2) or RNA3 (ΔRNA3). Evident retention of GFP fluorescence at 6 dpi was observed only in tissues infiltrated with the complete ToFBV genome (data not shown). This result suggests that RNA2 and RNA3 may contribute to the proper function of the RNA4-encoded ORF1 suppressor, potentially through interactions with other viral components during infection.

Overall, these results indicate that RNA4 plays multiple roles during ToFBV infection, contributing both to viral movement and to the suppression of host RNA silencing defences.

## Conclusion

In this study, we developed and characterized an agroinfiltration-based system for the delivery of the tomato fruit blotch virus (ToFBV) genome in *N. benthamiana* and tomato, providing new insights into the replication, protein function, and expression capacity of this member of the *Blunervirus* family. Our results demonstrate that the constructed agroclones are capable of initiating local infection in *N. benthamiana* and tomato, as evidenced by the accumulation of ssRNA1(−), a hallmark of active viral replication. Importantly, replication was strictly dependent on the presence of RNA2, confirming the central role of the viral RNA-dependent RNA polymerase in initiating replication in planta.

Systemic accumulation was detected in *N. benthamiana* at the later time point (>10 dpi), demonstrating that the developed agroclone system is capable of supporting long-distance viral movement within the host plant. ToFBV RNA accumulation in systemic leaves decreased at 15 dpi compared with earlier time points, suggesting that *N. benthamiana* may not provide an optimal host environment for sustained ToFBV accumulation in systemic leaves. Further experiments at additional time points, will be required to determine whether the observed decline is consistent and to better characterize the dynamics of ToFBV systemic infection.

A key outcome of this work is the functional validation of the ToFBV RNA3 and RNA4 segments. RNA3-derived constructs supported robust heterologous protein expression, and GFP insertion experiments revealed that ORF5 disruption provides a more favourable genomic context for expression than ORF2 disruption. This establishes ToFBV as a promising scaffold for the development of plant viral expression vectors, including in its natural host, where local replication of GFP-expressing constructs was demonstrated for the first time in an engineered *Blunervirus* system.

Structural and functional analyses further clarified the roles of RNA4-encoded proteins. ORF2 was identified as a bona fide movement protein, capable of complementing cell-to-cell movement in heterologous viral systems, consistent with AlphaFold and Foldseek predictions showing similarity to known viral movement proteins. In contrast, ORF1 showed no direct role in movement but displayed PTGS suppressor activity, suggesting a regulatory function in counteracting host antiviral defense mechanisms.

Finally, structural modelling of RNA3 ORFan proteins revealed both conserved and highly divergent features within the *Kitaviridae*, supporting a mixed evolutionary landscape where certain proteins (such as SP24-like ORF3) are structurally conserved, while others remain structurally and functionally uncharacterized.

Overall, our findings provide a first functional framework for ToFBV gene expression and protein activity, demonstrating that the virus encodes a minimal set of determinants sufficient for local replication, movement, and silencing suppression. *N. benthamiana* supports initial infection and sustained systemic spread of the virus. In tomato, local ToFBV replication was demonstrated by the accumulation of ssRNA1(-) and GFP, providing evidence of active viral replication in infected tissues. Further studies will focus on achieving systemic infection in tomato, the natural host of ToFBV.

These results not only deepen our understanding of ToFBV biology but also provide a foundation for future studies on kitavirus replication mechanisms and the development of novel plant viral vectors. Moreover, the biotechnological tools developed in this study may facilitate the identification of host factors involved in susceptibility and resistance, as well as support the development of diagnostic and control strategies to mitigate the impact of ToFBV and potentially other kitavirids in economically important crops.

## Acknowledgements

The authors performed experiments at the Imaging Platform of the Department of Translational and Molecular Medicine of the University of Brescia.

## CRediT

Niccolò Miotti: Conceptualization (equal), Writing – original draft (lead), Formal analysis (equal), Writing – review and editing (lead), Visualization (lead), Investigation (lead); Federica Bono: Investigation (equal); Giulia Golzi: Investigation (equal); Claudio Ratti: Resources (equal); Paola Casati: Writing – Review & Editing (equal); Massimo Turina: Conceptualization (lead), Formal analysis (equal), Writing – review and editing (equal), Investigation (equal), Resources (equal); Marina Ciuffo: Funding Acquisition (lead), Resources (equal).

## Data availability statement

The data that supports the findings of this study are available in the supplementary material of this article.

## SUPPLEMENTARY INFORMATIONS

**Supplementary Table 1:**
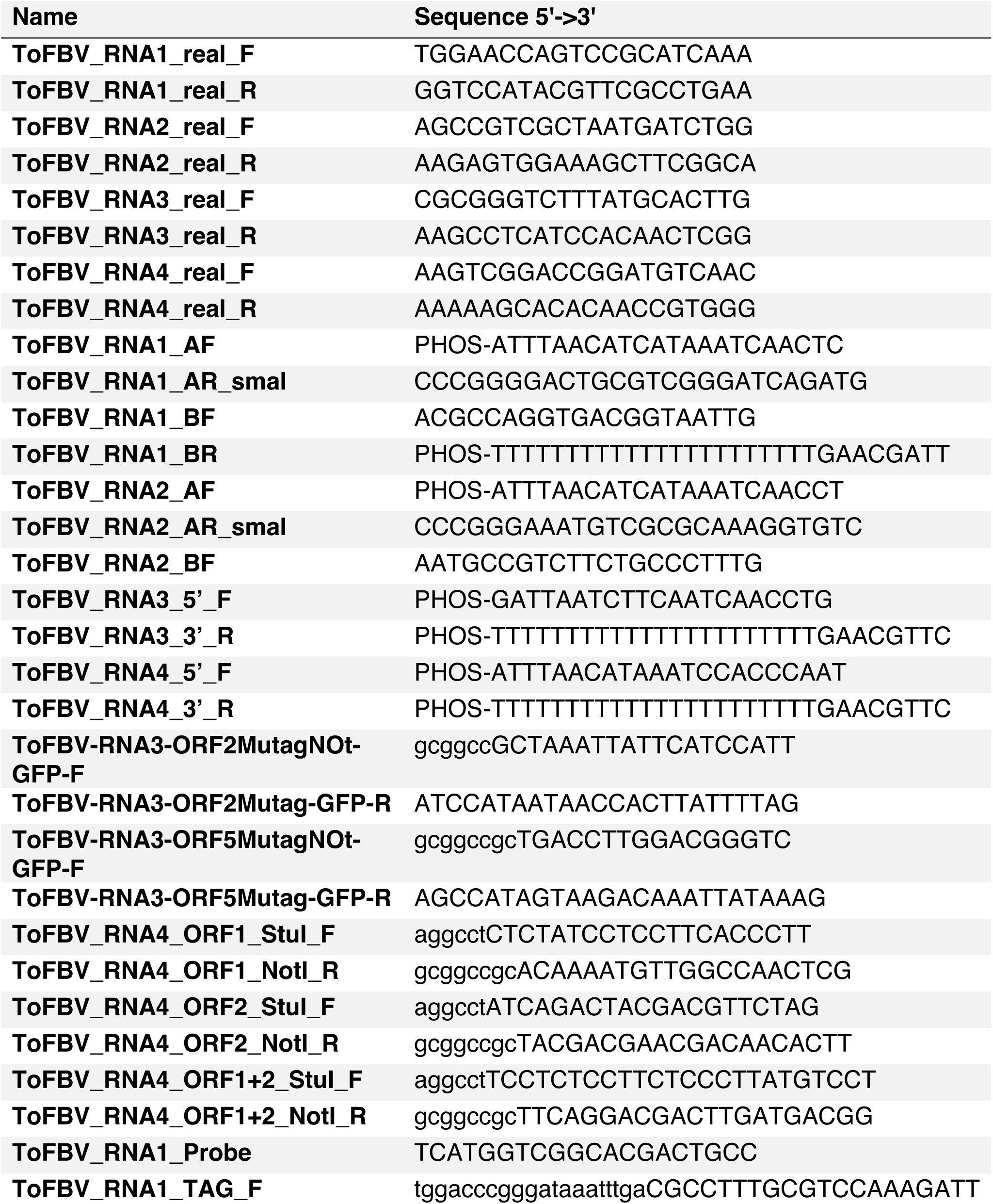
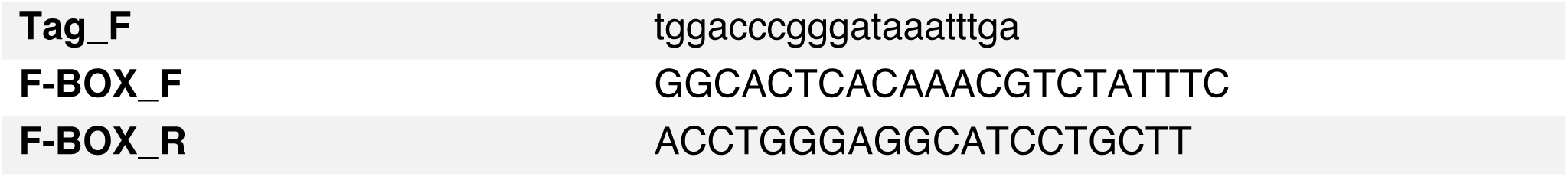
Primers used in this study are reported in the 5′→3′ orientation. Primers containing “real” in their name were used for the detection of viral fragments by RT-qPCR. Primers carrying a phosphorylated 5′ and are indicated by the prefix “PHOS” in the sequence. Lowercase letters represent restriction enzyme recognition sites or tag sequence that were added to the primer sequences.

**Supplementary table 2:** Comparative analysis of nucleotide substitutions and coding variations between Open Reading Frames (ORFs). Summary of identified mutations within the analyzed coding sequences, including codon positions, nucleotide coordinates, corresponding reference and mutant amino acids, and their functional classification (synonymous vs. non-synonymous).

| Name | AA-position | nts-position | Reference | Mutant | AA-Ref | AA-Mut | Type |
| --- | --- | --- | --- | --- | --- | --- | --- |
| ORF1_R NA1 | 6 | 16-18 | GCC | GCT | A | A | Synonymous |
| ORF1_R NA1 | 75 | 223-225 | TCC | TCT | S | S | Synonymous |
| ORF1_R NA1 | 102 | 304-306 | GTT | GTC | V | V | Synonymous |
| ORF1_R NA1 | 300 | 898-900 | TTC | TTT | F | F | Synonymous |
| ORF1_R NA1 | 386 | 1156-1158 | GAT | GAC | D | D | Synonymous |
| ORF1_R NA1 | 509 | 1525-1527 | TTT | TTC | F | F | Synonymous |
| ORF1_R NA1 | 515 | 1543-1545 | AAC | AAT | N | N | Synonymous |
| ORF1_R NA1 | 542 | 1624-1626 | GCT | GCC | A | A | Synonymous |
| ORF1_R NA1 | 603 | 1807-1809 | GTC | GTT | V | V | Synonymous |
| ORF1_R NA1 | 777 | 2329-2331 | CGT | CGC | R | R | Synonymous |
| ORF1_R NA1 | 782 | 2344-2346 | CTA | CTG | L | L | Synonymous |
| ORF1_R NA1 | 802 | 2404-2406 | TCT | TCC | S | S | Synonymous |
| ORF1_R NA1 | 805 | 2413-2415 | ATA | TTA | I | L | **Non-Synonymous** |
| ORF1_R NA1 | 873 | 2617-2619 | TTT | TTC | F | F | Synonymous |
| ORF1_R NA1 | 875 | 2623-2625 | TAT | TAC | Y | Y | Synonymous |
| ORF1_R NA1 | 909 | 2725-2727 | TCG | TTG | S | L | **Non-Synonymous** |
| ORF1_R NA1 | 942 | 2824-2826 | AAA | AAG | K | K | Synonymous |
| ORF1_R NA1 | 948 | 2842-2844 | TCT | TCC | S | S | Synonymous |
| ORF1_R<br>NA1 | 1320 | 3958-3960 | CAA | CTA | Q | L | **Non-Synonymous** |
| ORF1_R<br>NA1 | 1323 | 3967-3969 | CCT | GCT | P | A | **Non-Synonymous** |
| ORF1_R<br>NA1 | 1415 | 4243-4245 | AGA | AAA | R | K | **Non-Synonymous** |
| ORF1_R<br>NA1 | 1427 | 4279-4281 | TTT | TTC | F | F | Synonymous |
| ORF1_R<br>NA1 | 1432 | 4294-4296 | TCT | GCT | S | A | **Non-Synonymous** |
| ORF1_R<br>NA1 | 1433 | 4297-4299 | GAC | GAT | D | D | Synonymous |
| ORF1_R<br>NA1 | 1466 | 4396-4398 | GTG | GTA | V | V | Synonymous |
| ORF1_R<br>NA1 | 1495 | 4483-4485 | GAG | GAA | E | E | Synonymous |
| ORF1_R<br>NA1 | 1496 | 4486-4488 | CTC | CTT | L | L | Synonymous |
| ORF1_R<br>NA1 | 1500 | 4498-4500 | ACA | ACT | T | T | Synonymous |
| ORF1_R<br>NA1 | 1508 | 4522-4524 | ATC | ATT | I | I | Synonymous |
| ORF1_R<br>NA1 | 1513 | 4537-4539 | AGT | AGC | S | S | Synonymous |
| ORF1_R<br>NA1 | 1605 | 4813-4815 | AGT | AGC | S | S | Synonymous |
| ORF1_R<br>NA1 | 1632 | 4894-4896 | TTG | TTA | L | L | Synonymous |
| ORF1_R<br>NA1 | 1633 | 4897-4899 | GTC | GTT | V | V | Synonymous |
| ORF1_R<br>NA1 | 1643 | 4927-4929 | ATC | ATA | I | I | Synonymous |
| ORF1_R<br>NA1 | 1687 | 5059-5061 | TCC | TCT | S | S | Synonymous |
| ORF1_R<br>NA1 | 1689 | 5065-5067 | CTG | TTG | L | L | Synonymous |
| ORF1_R<br>NA1 | 1695 | 5083-5085 | AAG | AGG | K | R | **Non-Synonymous** |
| ORF1_R<br>NA1 | 1755 | 5263-5265 | GCT | GCA | A | A | Synonymous |
| ORF1_R<br>NA1 | 1760 | 5278-5280 | ACT | ACC | T | T | Synonymous |
| ORF1_R<br>NA1 | 1820 | 5458-5460 | AAA | AGA | K | R | **Non-Synonymous** |
| ORF1_R<br>NA2 | 7 | 19-21 | GAT | GAC | D | D | Synonymous |
| ORF1_R<br>NA2 | 16 | 46-48 | AAC | ACC | N | T | **Non-Synonymous** |
| ORF1_R<br>NA2 | 18 | 52-54 | CTT | CCT | L | P | **Non-Synonymous** |
| ORF1_R<br>NA2 | 41 | 121-123 | AAA | AAT | K | N | **Non-Synonymous** |
| ORF1_R<br>NA2 | 100 | 298-300 | AAA | AAG | K | K | Synonymous |
| ORF1_R<br>NA2 | 105 | 313-315 | ACC | ACT | T | T | Synonymous |
| ORF1_R<br>NA2 | 152 | 454-456 | TCT | TCC | S | S | Synonymous |
| ORF1_R<br>NA2 | 168 | 502-504 | CCG | CCA | P | P | Synonymous |
| ORF1_R<br>NA2 | 174 | 520-522 | AAG | AAA | K | K | Synonymous |
| ORF1_R<br>NA2 | 178 | 532-534 | CCT | CCC | P | P | Synonymous |
| ORF1_R<br>NA2 | 182 | 544-546 | CGG | CGA | R | R | Synonymous |
| ORF1_R<br>NA2 | 187 | 559-561 | GTC | GTT | V | V | Synonymous |
| ORF1_R<br>NA2 | 192 | 574-576 | CCC | CCT | P | P | Synonymous |
| ORF1_R<br>NA2 | 519 | 1555-1557 | GCG | GCA | A | A | Synonymous |
| ORF1_R<br>NA2 | 563 | 1687-1689 | CGC | CGT | R | R | Synonymous |
| ORF1_R<br>NA2 | 568 | 1702-1704 | CTT | CTC | L | L | Synonymous |
| ORF1_R<br>NA2 | 578 | 1732-1734 | GTT | ATC | V | I | **Non-Synonymous** |
| ORF1_R<br>NA2 | 585 | 1753-1755 | AAT | AAC | N | N | Synonymous |
| ORF1_R<br>NA2 | 593 | 1777-1779 | GCT | GTT | A | V | **Non-Synonymous** |
| ORF1_R<br>NA2 | 827 | 2479-2481 | ATT | ATC | I | I | Synonymous |
| ORF1_R<br>NA2 | 953 | 2857-2859 | GTT | GTC | V | V | Synonymous |
| ORF1_R<br>NA2 | 1114 | 3340-3342 | GAG | GAA | E | E | Synonymous |
| ORF5_R<br>NA3 | 179 | 535-537 | CCA | CAA | P | Q | **Non-Synonymous** |
| ORF2_R<br>NA4 | 194 | 568-570 | GCC | GCT | A | A | Synonymous |
| ORF2_R<br>NA4 | 308 | 922-924 | CCT | CCC | P | P | Synony<br>mous |

**Supplementary Table 3.** RT-qPCR detection of ToFBV RNAs in tomato and *Nicotiana benthamiana*. The table reports the Ct values and standard deviations (StdDev Ct) obtained for the detection of ToFBV RNAs, and negative-sense RNA1 [ssRNA1(−)] in local and systemic leaf samples. The indicated detection chemistry (SYBR Green or TaqMan) is reported for each assay. Samples include locally infiltrated tomato leaves, systemically infected *N. benthamiana* used as a control, and systemic leaves of *N. benthamiana* analyzed at 10 and 15 dpi. At 10 dpi, ssRNA1(−) accumulation was compared between plants infiltrated with the complete ToFBV (compl) construct and the ΔRNA2 combination. Ct values are reported together with their corresponding standard deviations.

| *Sample Name* | *Detector* | *Chemistry* | *Ct* | *StdDev*<br>*v Ct* |
| --- | --- | --- | --- | --- |
| ΔRNA2 Tom-cDNA | ToFBV_RNA1 | Taq-Man | 35.8625 | 0.291-999.9 |
| ΔRNA2 Tom-cDNA | ToFBV_RNA1 | Taq-Man | 35.4516 | 0.291-999.9 |
| ΔRNA2 Tom-DNAse | ToFBV_RNA1 | Taq-Man | Undetermined | |
| ΔRNA2 Tom-DNAse | ToFBV_RNA1 | Taq-Man | Undetermined | |
| ΔRNA2 TomTNA | ToFBV_RNA1 | Taq-Man | 32.5182 | 0.283-999.9 |
| ΔRNA2 TomTNA | ToFBV_RNA1 | Taq-Man | 32.1179 | 0.283-999.9 |
| Compl-Tom-cDNA | ToFBV_RNA1 | Taq-Man | 31.9527 | 0.087<br>7-999.9 |
| Compl-Tom-cDNA | ToFBV_RNA1 | Taq-Man | 31.8287 | 0.087<br>7-999.9 |
| Compl-Tom-DNAse | ToFBV_RNA1 | Taq-Man | Undetermined |  |
| Compl-Tom-DNAse | ToFBV_RNA1 | Taq-Man | Undetermined |  |
| Compl-Tom-TNA | ToFBV_RNA1 | Taq-Man | 33.518 | 0.344-999.9 |
| Compl-Tom-TNA | ToFBV_RNA1 | Taq-Man | 33.0308 | 0.344-999.9 |
| Syst benth Compl | ToFBV_RNA1 | Taq-Man | 27.7334 | 0.111-999.9 |
| Syst benth Compl | ToFBV_RNA1 | Taq-Man | 27.5771 | 0.111-999.9 |

**Detection of RNA4 locally in tomato, and as control, *N. benthamiana* systemically infected**
| *Sample Name* | *Detector* | *Chemistry* | *Ct* | *StdDev*<br>*v Ct* |
| --- | --- | --- | --- | --- |
| ΔRNA2 Tom-cDNA | ToFBV RNA 4 | SYBER-Green | 29.8089 | 0.217-999.9 |
| ΔRNA2 Tom-cDNA | ToFBV RNA 4 | SYBER-Green | 30.1164 | 0.217-999.9 |
| ΔRNA2 Tom-DNAse | ToFBV RNA 4 | SYBER-Green | 35.0764 | 1.34-999.9 |
| ΔRNA2 Tom-DNAse | ToFBV RNA 4 | SYBER-Green | 36.9657 | 1.34-999.9 |
| ΔRNA2 TomTNA | ToFBV RNA 4 | SYBER-Green | 26.3754 | 0.0964-999.9 |
| ΔRNA2 TomTNA | ToFBV RNA 4 | SYBER-Green | 26.2391 | 0.0964-999.9 |
| Compl-Tom-cDNA | ToFBV RNA 4 | SYBER-Green | 26.2253 | 0.0289-999.9 |
| Compl-Tom-cDNA | ToFBV RNA 4 | SYBER-Green | 26.1844 | 0.0289-999.9 |
| Compl-Tom-DNAse | ToFBV RNA 4 | SYBER-Green | Undetermined |  |
| Compl-Tom-DNAse | ToFBV RNA 4 | SYBER-Green | 35.4505 | -999.9 |
| Compl-Tom-TNA | ToFBV RNA 4 | SYBER-Green | 26.9468 | 0.206-999.9 |
| Compl-Tom-TNA | ToFBV RNA 4 | SYBER-Green | 27.2381 | 0.206-999.9 |
| NTC | ToFBV RNA 4 | SYBER-Green | 35.9126 | -999.9 |
| Syst benth Compl | ToFBV RNA 4 | SYBER-Green | 19.1284 | 0.218-999.9 |
| Syst benth Compl | ToFBV RNA 4 | SYBER-Green | 19.4362 | 0.218-999.9 |

Detection of ssRNA1(-) locally in tomato
| *Sample Name* | *Detector* | *Chemestry* | *Ct* | *StdDev Ct* |
| --- | --- | --- | --- | --- |
| ΔRNA2 Tom-cDNA | ToFBV_ssRNA1(-)_tag | Taq-Man | 40.00 | 0.248 |
| ΔRNA2 Tom-cDNA | ToFBV_ssRNA1(-)_tag | Taq-Man | 38.6628 | 0.248-999.9 |
| ΔRNA2 Tom-cDNA | ToFBV_ssRNA1(-)_tag | Taq-Man | 38.3127 | 0.248-999.9 |
| Compl-Tom-cDNA | ToFBV_ssRNA1(-)_tag | Taq-Man | 36.313 | 0.379-999.9 |
| Compl-Tom-cDNA | ToFBV_ssRNA1(-)_tag | Taq-Man | 37.0637 | 0.379-999.9 |
| Compl-Tom-cDNA | ToFBV_ssRNA1(-)_tag | Taq-Man | 36.7738 | 0.379-999.9 |
| TNAdnaseTom-2 | ToFBV_ssRNA1(-)_tag | Taq-Man | Undetermined |  |
| TNAdnaseTom-2 | ToFBV_ssRNA1(-)_tag | Taq-Man | Undetermined |  |
| TNAdnaseTom-2 | ToFBV_ssRNA1(-)_tag | Taq-Man | Undetermined |  |
| TNAdnasPomCompl | ToFBV_ssRNA1(-)_tag | Taq-Man | Undetermined |  |
| TNAdnasPomCompl | ToFBV_ssRNA1(-)_tag | Taq-Man | Undetermined |  |
| TNAdnasPomCompl | ToFBV_ssRNA1(-)_tag | Taq-Man | Undetermined |  |

***N. Benthamiana* systemic: comparison of complete mix and ΔRNA2 mix. at 10 dpi**
| *Sample Name* | *Detector* | *Chemestry* | *Ct* | *StdDe*<br>*v Ct* |
| --- | --- | --- | --- | --- |
| ToFBV complete | ToFBV_RNA1 | Taq-Man | 26.5509 | 0.060<br>4-999.9 |
| ToFBV complete | ToFBV_RNA1 | Taq-Man | 26.6363 | 0.060<br>4-999.9 |
| ToFBV complete | ToFBV_RNA2 | SYBER-Green | 26.1076 | 0.744-999.9 |
| ToFBV complete | ToFBV_RNA2 | SYBER-Green | 25.0561 | 0.744-999.9 |
| ToFBV complete | ToFBV_RNA3 | SYBER-Green | 21.9225 | 0.050<br>1-999.9 |
| ToFBV complete | ToFBV_RNA3 | SYBER-Green | 21.9933 | 0.050<br>1-999.9 |
| ToFBV complete | ToFBV RNA 4 | SYBER-Green | 22.2239 | 2.02-999.9 |
| ToFBV complete | ToFBV RNA 4 | SYBER-Green | 19.3656 | 2.02-999.9 |
| ToFBV complete | ToFBV_RNA1 | Taq-Man | Undetermined |  |
| compl-noRT | ToFBV_RNA1 | Taq-Man | Undetermined |  |
| compl-noRT | ToFBV_RNA2 | SYBER-Green | Undetermined |  |
| compl-noRT | ToFBV_RNA2 | SYBER-Green | 35.5739 | -999.9 |
| compl-noRT | ToFBV_RNA3 | SYBER-Green | 33.1965 | -999.9 |
| compl-noRT | ToFBV_RNA3 | SYBER-Green | Undetermined |  |
| compl-noRT | ToFBV RNA 4 | SYBER-Green | Undetermined |  |
| compl-noRT | ToFBV RNA 4 | SYBER-Green | 34.3581 | -999.9 |
| ΔRNA2 | ToFBV_RNA1 | Taq-Man | 38.5625 | -999.9 |
| ΔRNA2 | ToFBV_RNA1 | Taq-Man | Undetermined |  |
| ΔRNA2 | ToFBV_RNA2 | SYBER-Green | 32.1422 | 1.18-999.9 |
| ΔRNA2 | ToFBV_RNA2 | SYBER-Green | 30.4703 | 1.18-999.9 |
| ΔRNA2 | ToFBV_RNA3 | SYBER-Green | 32.4578 | 1.3-999.9 |
| ΔRNA2 | ToFBV_RNA3 | SYBER-Green | 30.6181 | 1.3-999.9 |
| ΔRNA2 | ToFBV RNA 4 | SYBER-Green | 32.427 | 0.869-999.9 |
| ΔRNA2 | ToFBV RNA 4 | SYBER-Green | 31.1975 | 0.869-999.9 |

**Detection of all ToFBV RNAs in systemic leaves of *N. benthamiana* at 15 dpi**
| *Sample Name* | *Detector* | *Chemestry* | *Ct* | *StdDe*<br>*v Ct* |
| --- | --- | --- | --- | --- |
| 2 | RNA1 | SYBER-Green | 30,35713959 | 0,375<br>46300 |
| 2 | RNA1 | SYBER-Green | 29,82615471 | 0,375<br>46300 |
| 2 | RNA2 | SYBER-Green | 33,72304535 | 0,342<br>82583 |
| 2 | RNA2 | SYBER-Green | 34,2078743 | 0,342<br>82583 |
| 2 | RNA3 | SYBER-Green | 30,02892685 | 0,251<br>26534 |
| 2 | RNA3 | SYBER-Green | 29,67358398 | 0,251<br>26534 |
| 2 | RNA4 | SYBER-Green | 28,16424179 | 3,767<br>51017 |
| 2 | RNA4 | SYBER-Green | 33,49230576 | 3,767<br>51017 |
| 4 | RNA1 | SYBER-Green | 30,59057426 | 1,672<br>55556 |
| 4 | RNA1 | SYBER-Green | 28,22522354 | 1,672<br>55556 |
| 4 | RNA2 | SYBER-Green | 32,2081604 | 0,758<br>47333 |
| 4 | RNA2 | SYBER-Green | 33,28080368 | 0,758<br>47333 |
| 4 | RNA3 | SYBER-Green | 29,77882385 | 0,079<br>24281 |
| 4 | RNA3 | SYBER-Green | 29,89089012 | 0,079<br>24281 |
| 4 | RNA4 | SYBER-Green | 29,2944622 | 0,465<br>73144 |
| 4 | RNA4 | SYBER-Green | 29,95310593 | 0,465<br>73144 |
| 11 | RNA1 | SYBER-Green | 30,99876022 | 0,475<br>17907 |
| 11 | RNA1 | SYBER-Green | 30,32675552 | 0,475<br>17907 |
| 11 | RNA2 | SYBER-Green | 33,22219086 | 0,535<br>61431 |
| 11 | RNA2 | SYBER-Green | 33,97966385 | 0,535<br>61431 |
| 11 | RNA3 | SYBER-Green | 28,41783905 | 0,258<br>09246 |
| 11 | RNA3 | SYBER-Green | 28,78283691 | 0,258<br>09246 |
| 11 | RNA4 | SYBER-Green | 30,68820000 | 0,067<br>40591 |
| 11 | RNA4 | SYBER-Green | 30,59287364 | 0,067<br>40591 |
| 13 | RNA1 | SYBER-Green | 30,19444275 | 0,895<br>99883 |
| 13 | RNA1 | SYBER-Green | 31,46157646 | 0,895<br>99883 |
| 13 | RNA2 | SYBER-Green | 33,48916626 | 0,854<br>09610 |
| 13 | RNA2 | SYBER-Green | 34,69704056 | 0,854<br>09610 |
| 13 | RNA3 | SYBER-Green | 29,67327309 | 0,131<br>89603 |
| 13 | RNA3 | SYBER-Green | 29,85980225 | 0,131<br>89603 |
| 13 | RNA4 | SYBER-Green | 31,60682678 | 0,077<br>40489 |
| 13 | RNA4 | SYBER-Green | 31,71629383 | 0,077<br>40489 |

**Detection of ToFBV RNA1 in systemic leaves of *N. benthamiana* at 15 dpi**
| *Sample Name* | *Detector* | *Chemestry* | *Ct* | *StdDev*<br>*v Ct* |
| --- | --- | --- | --- | --- |
| 1 | RNA1 | SYBER-Green | 30,50725555 | 0,769<br>18876 |
| 1 | RNA1 | SYBER-Green | 29,41945839 | 0,769<br>18876 |
| 2 | RNA1 | SYBER-Green | 30,35713959 | 0,375<br>46300 |
| 2 | RNA1 | SYBER-Green | 29,82615471 | 0,375<br>46300 |
| 3 | RNA1 | SYBER-Green | 29,79285431 | 0,057<br>37905 |
| 3 | RNA1 | SYBER-Green | 29,71170807 | 0,057<br>37905 |
| 4 | RNA1 | SYBER-Green | 30,59057426 | 1,672<br>55556 |
| 4 | RNA1 | SYBER-Green | 28,22522354 | 1,672<br>55556 |
| 5 | RNA1 | SYBER-Green | Undetermined |  |
| 5 | RNA1 | SYBER-Green | Undetermined |  |
| 6 | RNA1 | SYBER-Green | 29,37669754 | 0,322<br>17320 |
| 6 | RNA1 | SYBER-Green | 29,83231926 | 0,322<br>17320 |
| 7 | RNA1 | SYBER-Green | 30,28031349 | 0,379<br>38747 |
| 7 | RNA1 | SYBER-Green | 30,81684841 | 0,379<br>38747 |
| 8 | RNA1 | SYBER-Green | Undetermined |  |
| 8 | RNA1 | SYBER-Green | Undetermined |  |
| 9 | RNA1 | SYBER-Green | 30,41542435 | 0,390<br>39310 |
| 9 | RNA1 | SYBER-Green | 29,86332512 | 0,390<br>39310 |
| 10 | RNA1 | SYBER-Green | Undetermined |  |
| 10 | RNA1 | SYBER-Green | Undetermined |  |
| 11 | RNA1 | SYBER-Green | 30,99876022 | 0,475<br>17907 |
| 11 | RNA1 | SYBER-Green | 30,32675552 | 0,475<br>17907 |
| 12 | RNA1 | SYBER-Green | 31,62657547 | 0,249<br>87817 |
| 12 | RNA1 | SYBER-Green | 31,27319436 | 0,249<br>87817 |
| 13 | RNA1 | SYBER-Green | 30,19444275 | 0,895<br>99884 |
| 13 | RNA1 | SYBER-Green | 31,46157646 | 0,895<br>99884 |
| 14 | RNA1 | SYBER-Green | 31,37827110 | 0,161<br>61327 |
| 14 | RNA1 | SYBER-Green | 31,60682678 | 0,161<br>61327 |

**Supplementary Figure 1:**
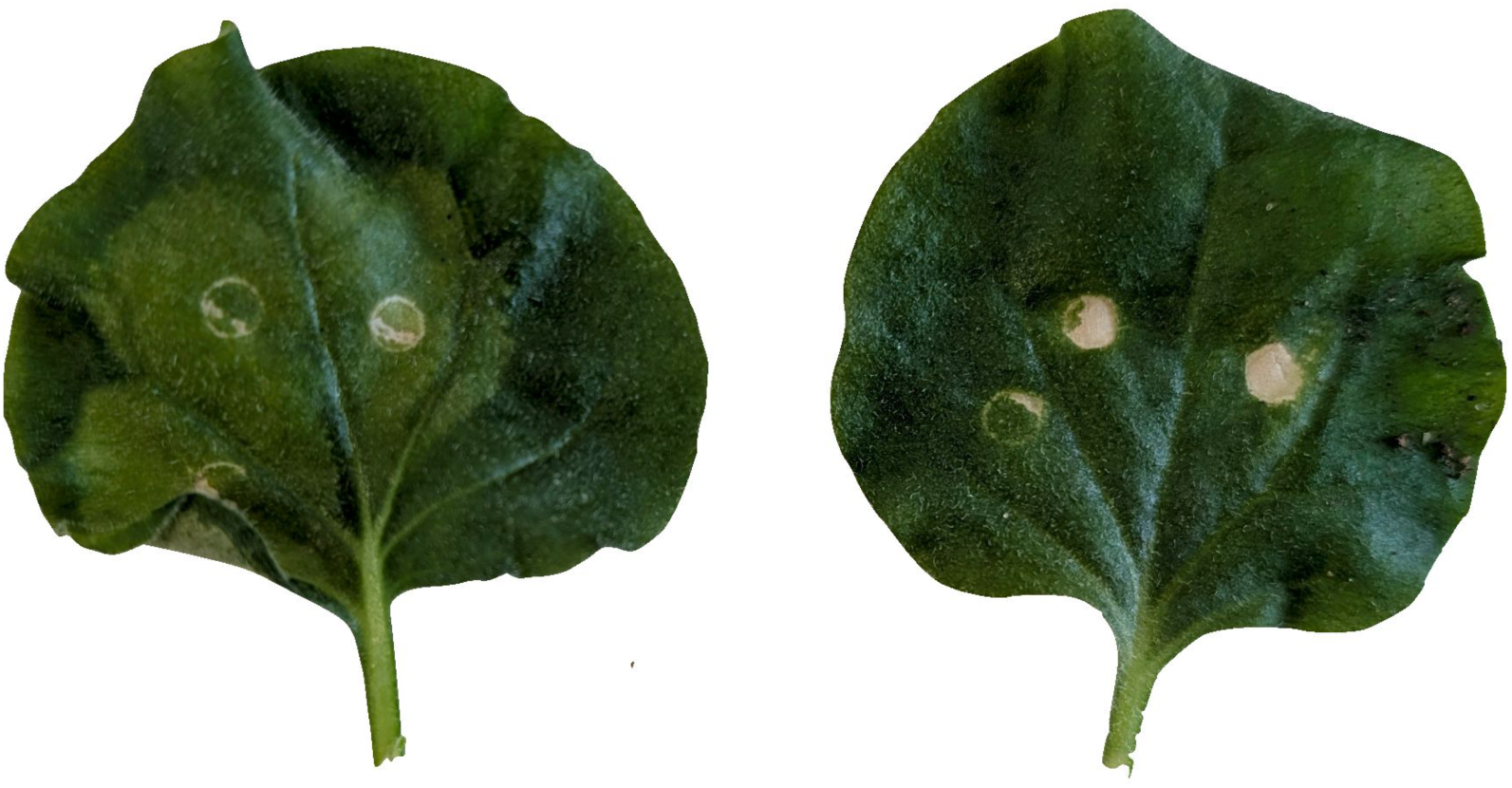
Patch areas of N. benthamiana leaves agroinfiltrated with the complete ToFBV construct (left) or ΔRNA2 (right) 5 days dpi. Localized discoloration is evident in tissues expressing the complete ToFBV, whereas no clear symptoms are observed in the ΔRNA2 treatment, apart from minor mechanical damage associated with the agroinfiltration procedure (needleless syringe).

**Supplementary Figure 2:**
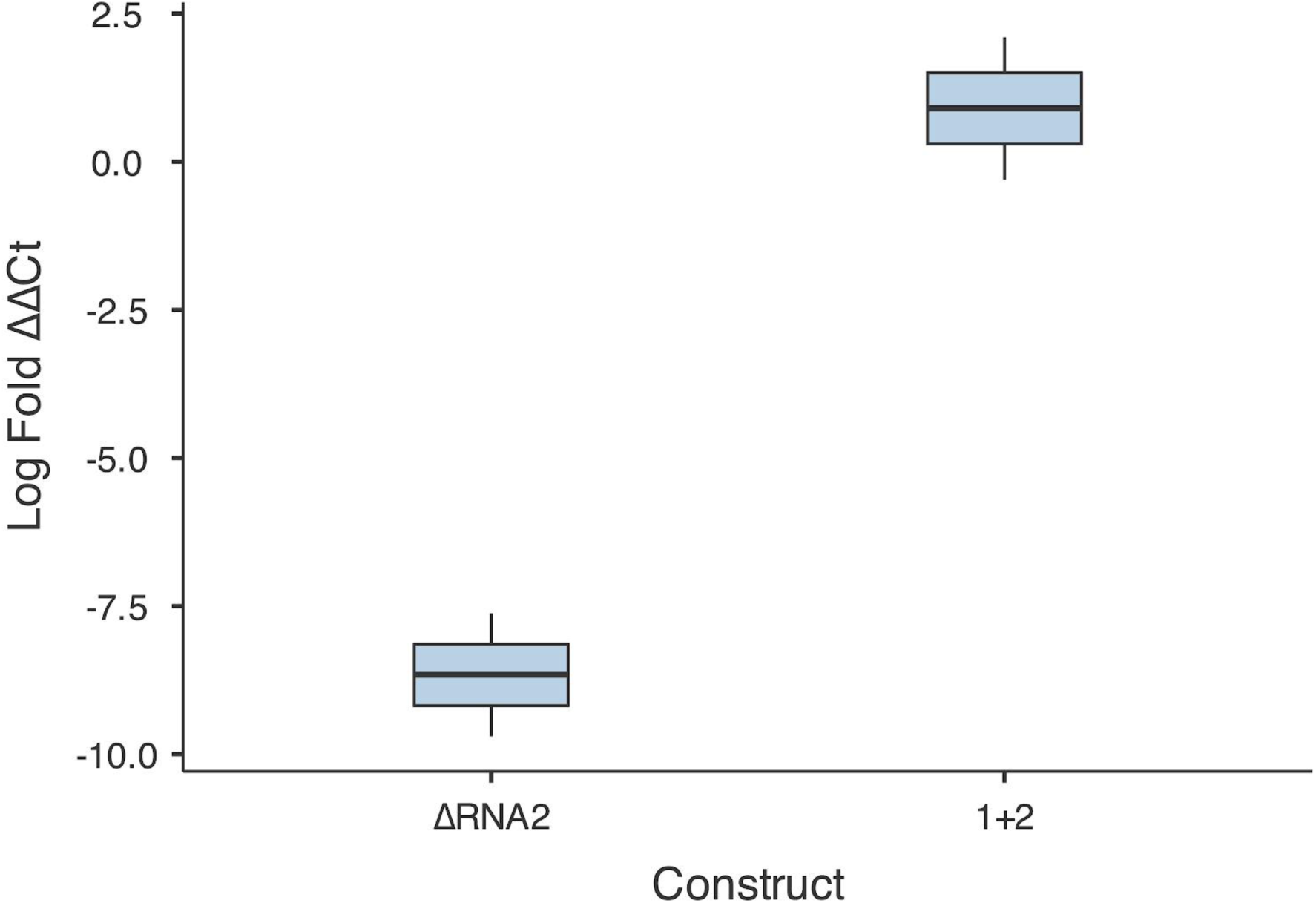
Relative accumulation of ssRNA1(-) in ΔRNA2 and 1+2 compared to ToFBV complete. Box plots display the log-transformed fold change (ΔΔCt) of ssRNA1(-) accumulation for the ΔRNA2 and 1+2 constructs. Expression levels for all samples are normalized against the F-BOX housekeeping gene and calculated relative to the ToFBV complete construct baseline (value equal to 0, not shown). The ΔRNA2 construct exhibits a profound reduction in ssRNA1(-) accumulation (median log fold change of approximately -8.5). Conversely, the 1+2 construct demonstrates accumulation levels comparable to or up to two-fold higher than the ToFBV complete control (median log fold change of approximately 1.0). Center lines represent the median, box limits indicate the 25th and 75th percentiles, and whiskers extend to the minimum and maximum data points.

**Supplementary Figure 3:**
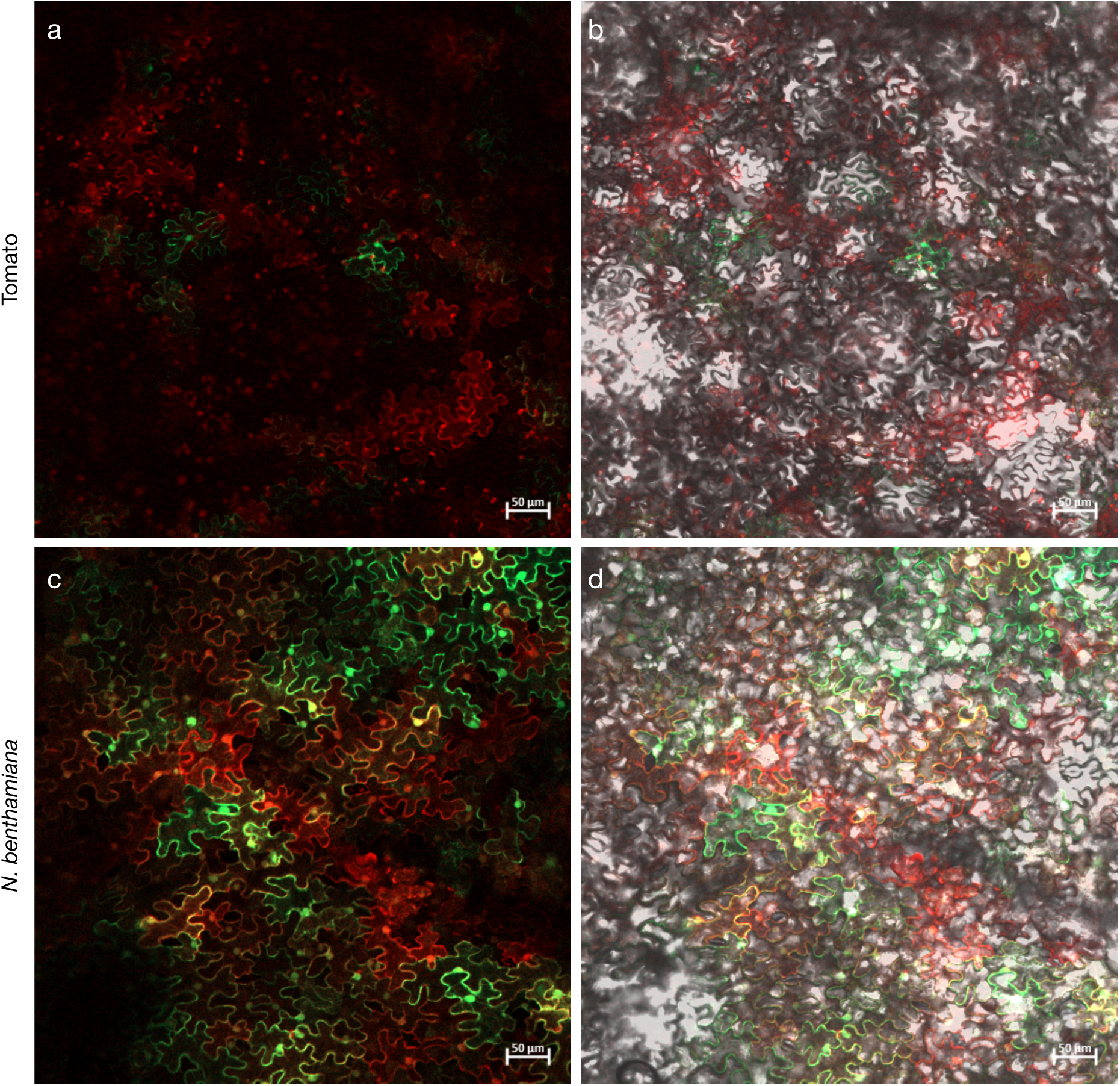
Comparison of transient co-transformation and co-expression efficiencies in different plant hosts. Confocal laser scanning microscopy of plant leaf epidermal cells following agroinfiltration with GFP and mCherry constructs. (a, b) Transient expression in tomato (*Solanum lycopersicum* cv Marmande) leaves, showing a low frequency of successful co-transformation and co-expression events. (a) Merged fluorescence channels; (b) Overlay of fluorescence and brightfield channels. (c, d) Transient expression in *N. benthamiana* leaves, displaying highly efficient, widespread co-transformation and co-expression across the epidermal cell layer. (c) Merged fluorescence channels; (d) Overlay of fluorescence and brightfield channels. The distinct delivery efficiencies observed between these hosts suggest significant host-dependent limitations in the uptake and expression of transient multi-construct systems, highlighting potential bottlenecks when attempting to replicate multipartite virus infections. Scale bars = 50 µm.

**Supplementary Figure 4:**
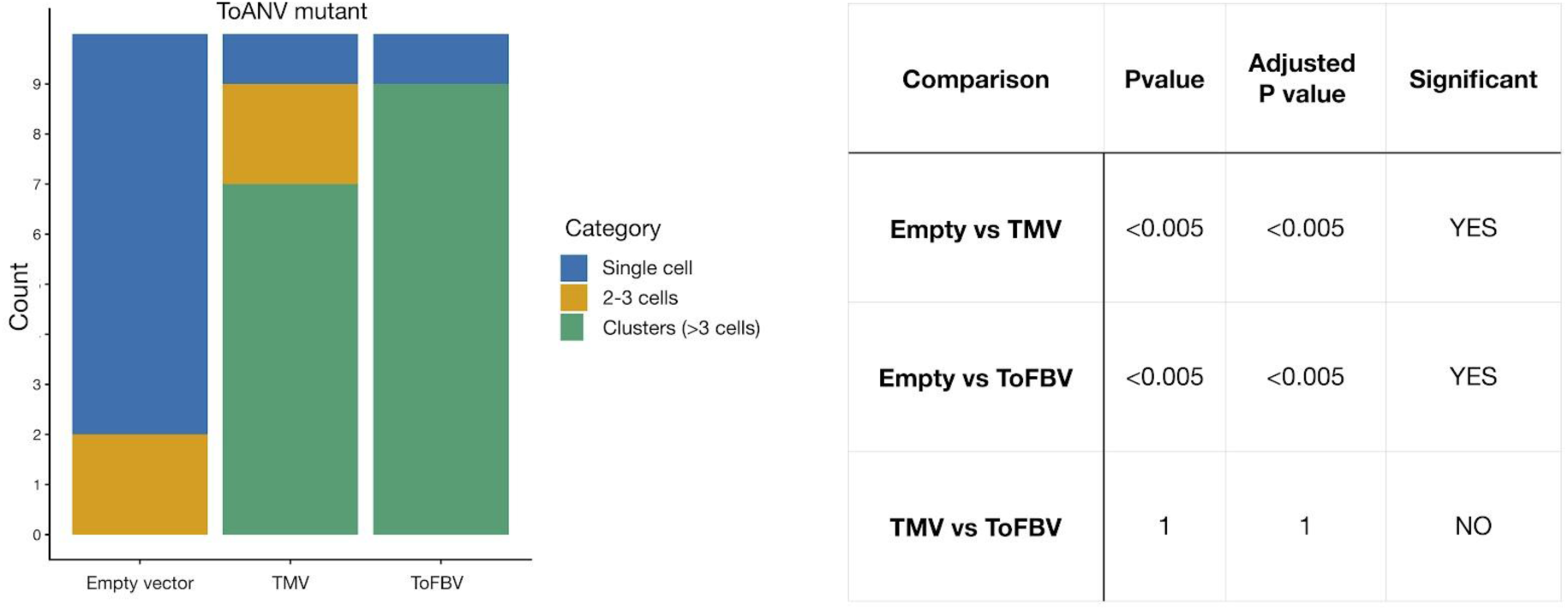
Functional complementation of an MP-deficient ToANV clone with heterologous movement proteins. Stacked bar graphs (tleft) categorize infection foci as single cells, 2–3 cell groups, or clusters (>3 cells) for the empty vector, TMV, and ToFBV constructs. The summary table (right) provides statistical comparisons (Adjusted P-values), indicating that ToFBV ORF2 significantly promotes viral spread compared to the negative control (p < 0.005) and performs at a level statistically comparable to the TMV MP in the ToANV system.

**Supplementary Figure 5:**
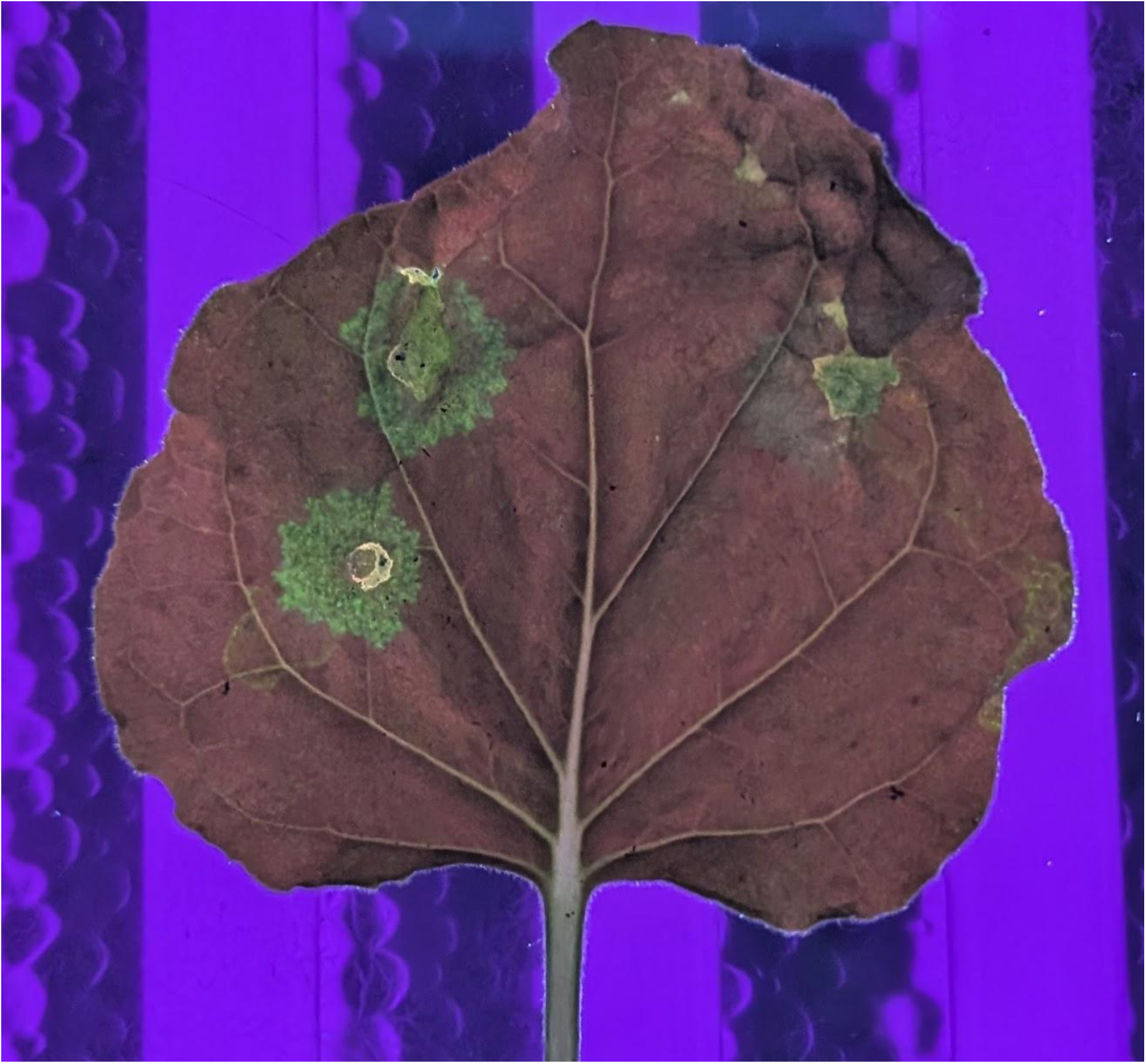
Visual assessment of PVX mutant movement protein complementation on a leaf under UV light. In vivo rescue of a movement-deficient PVX mutant expressing GFP, visualized under a UV transilluminator at 10 days dpi. The left side of the leaf was co-inoculated with the ToFBV MP, showing a significantly larger area of green fluorescent spread. The right side of the leaf was co-inoculated with the TMV MP, resulting in noticeably smaller, more restricted fluorescent foci.

**Supplementary Figure 6:**
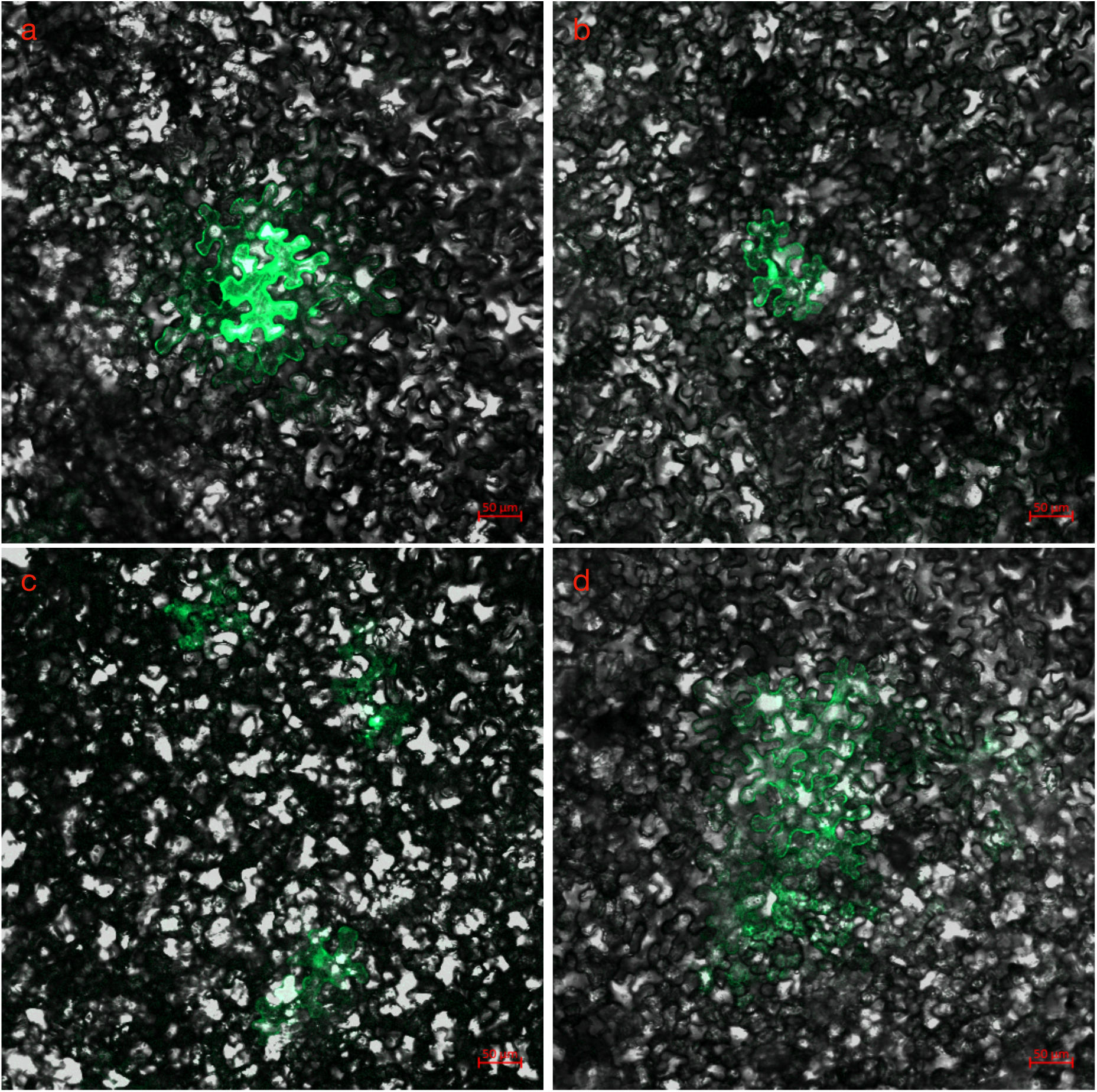
Confocal microscopic analysis of PVX movement protein complementation. Confocal laser scanning displaying the cell-to-cell movement of PVX mp-mutant expressing GFP in epidermal cells. Images represent high-resolution tracking of viral spread at 3 dpi. (a) Complementation with ToFBV ORF2, showing functional rescue of viral transport indicated by multicellular green fluorescent clusters. (b) Complementation with ToFBV ORF1, demonstrating a lack of movement with fluorescence restricted to a single cell. (c) Empty vector negative control, showing isolated single-cell fluorescence. (d) TMV MP positive control, demonstrating robust intercellular spread and large fluorescent foci. Scale bars = 50 µm.

